# Sensitivity and resolution improvement for in-vivo magnetic resonance current density imaging (MRCDI) of the human brain

**DOI:** 10.1101/2021.03.23.436558

**Authors:** Cihan Göksu, Klaus Scheffler, Fróði Gregersen, Hasan H. Eroğlu, Rahel Heule, Hartwig R. Siebner, Lars G. Hanson, Axel Thielscher

## Abstract

**Purpose:** Magnetic resonance current density imaging (MRCDI) combines MR brain imaging with the injection of time-varying weak currents (1-2 mA) to assess the current flow pattern in the brain. However, the utility of MRCDI is still hampered by low measurement sensitivity and poor image quality.

**Methods:** We recently introduced a multi-gradient-echo-based MRCDI approach that has the hitherto best documented efficiency. We now advanced our MRCDI approach in three directions and performed phantom and in-vivo human brain experiments for validation: First, we verified the importance of enhanced spoiling and optimize it for imaging of the human brain. Second, we improved the sensitivity and spatial resolution by using acquisition weighting. Third, we added navigators as a quality control measure for tracking physiological noise. Combining these advancements, we tested our optimized MRCDI method by using 1 mA transcranial electrical stimulation (TES) currents injected via two different electrode montages in five subjects.

**Results:** For a session duration of 4:20 min, the new MRCDI method was able to detect magnetic field changes caused by the TES current flow at a sensitivity level of 84 pT, representing in a twofold increase relative to our original method. Comparing both methods to current flow simulations based on personalized head models demonstrated a consistent increase in the coefficient of determination of Δ*R*^2^=0.12 for the current-induced magnetic fields and Δ*R*^2^=0.22 for the current flow reconstructions. Interestingly, some of the simulations still clearly deviated from the measurements despite of the strongly improved measurement quality. This suggests that MRCDI can reveal useful information for the improvement of head models used for current flow simulations.

**Conclusion:** The advanced method strongly improves the sensitivity and robustness of MRCDI and is an important step from proof-of-concept studies towards a broader application of MRCDI in clinical and basic neuroscience research.

## INTRODUCTION

Magnetic resonance current density imaging (MRCDI) and MR electrical impedance tomography (MREIT) are emerging modalities to measure the current flow of transcranial electric stimulation (TES) (1) and the ohmic tissue conductivities non-invasively in the human brain (2–12). They combine MRI with weak electrical currents of 1-2 mA baseline-to-peak, alternating at low frequencies (0-100 Hz; Fig. 1). The currents induce an alternating magnetic field in the brain, and its component Δ*B_z,c_* parallel to the magnetic field of the MR scanner causes tiny modulations of the MR phase. MRCDI and MREIT aim to measure these phase modulations to gain insight into the strength and spatial distribution of the current-induced magnetic field for informing reconstructions of the current flow and conductivity distributions. The approach has been successfully demonstrated in phantoms, animals and human limbs in-vivo (7,9,15–28), but it is particularly challenging for the human brain because the current-induced magnetic field stays below 1 to 2 nT in that case. This is caused by the low maximal current strength that is tolerable and safe (1-2 mA) (29) and the “shielding” effect of the low-conductive skull that results in part of the current to be shunted through the scalp.

**Figure 1.**
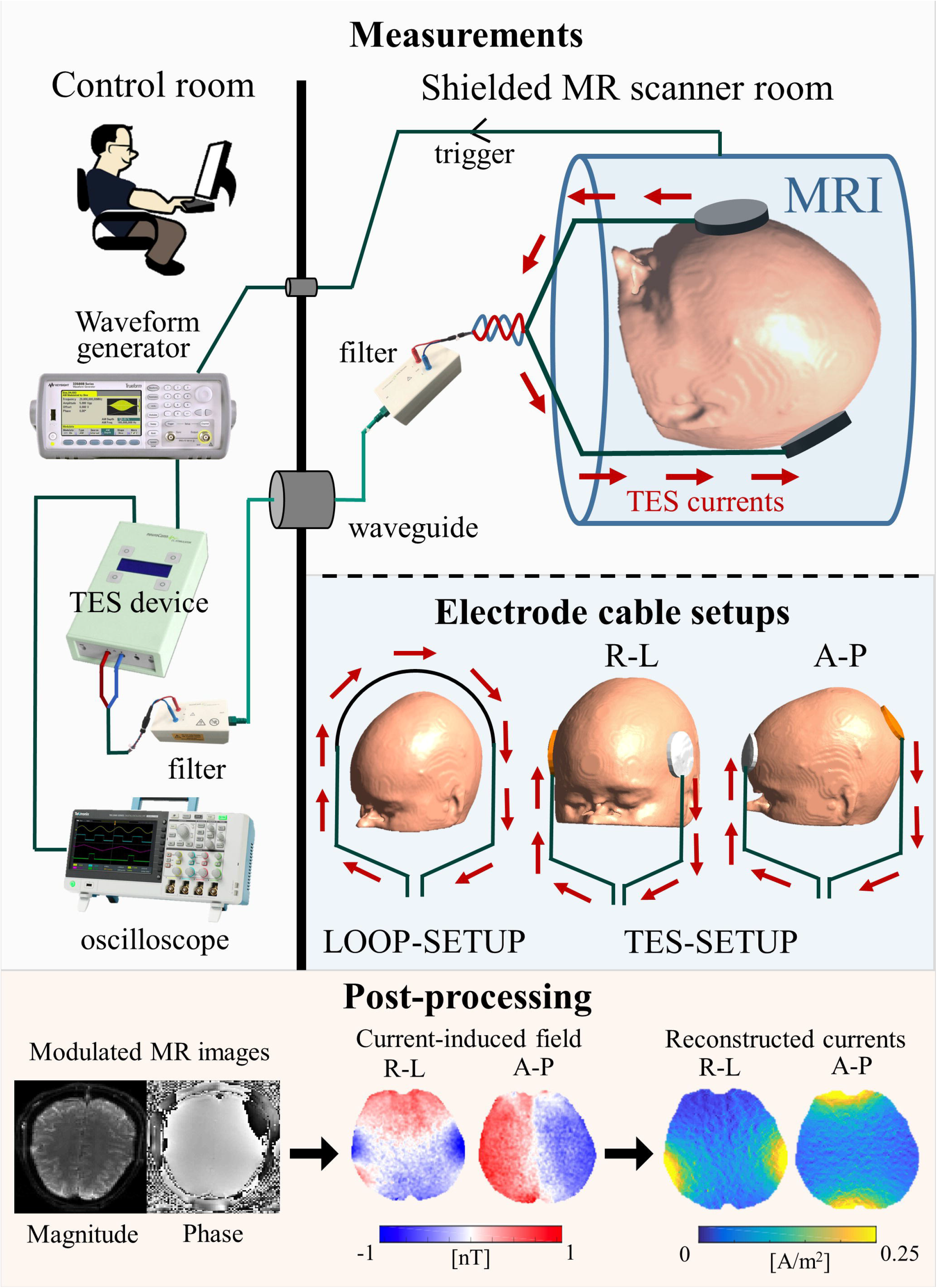
Experimental set-up for human in-vivo brain MRCDI. The MR scanner sends a trigger signal to an arbitrary waveform generator (33500B; Keysight Technologies, Santa Clara, CA, USA) in synchrony with the applied RF pulses. The generated voltage waveform determines the TES current strength, pulse-width, and polarity that are monitored with an oscilloscope. The voltage waveform is converted to electrical currents by a TES stimulator (DC-STIMULATOR MR, NeuroCare Group GmbH, Munich, Germany). The generated electrical currents are filtered from RF noise. Two different setups are used in this study. LOOP-SETUP: The currents are not injected into the head but pass through a cable loop placed around the head. TES-SETUP: The TES currents are injected via scalp electrodes placed according to the desired current injection profile; right-left (R-L) or anterior-posterior (A-P).

In prior proof-of-concept studies, we focused on optimization of the MR sequences (10,30) and on correction of the impact of stray magnetic fields caused by the currents in the cables connected to the TES electrodes (31). We demonstrated measurements of the current-induced magnetic field in the in-vivo human brain at a sensitivity of ~0.1 nT in a ~9 mins scan (10). While promising, these early results indicated the need for further improvements: First, additional sensitivity enhancements of the Δ*B_z,c_* measurements would be needed to obtain a good signal-to-noise ratio (SNR) in the reconstructed current flow and conductivity distributions. Second, the measurements were sensitive to physiological noise (e.g. due to subject movement), so that we occasionally had to exclude part of the data after visual inspection (10). This pointed towards the need for developing an independent marker of the quality of the Δ*B_z,c_* field measurements to amend the qualitative and subjective judgement. Third, in some Δ*B_z,c_* images, artefacts close to the cables remained visible even after correcting for the cable stray fields (e.g. Figure 4 in (31)). Further testing revealed that these artefacts were unlikely to stem mostly from inaccurate tracking of the cable paths, as initially thought, but rather pointed towards imperfect measurements.

In this study, we strongly improve our MRCDI approach based on gradient-echo imaging (10) to tackle the above challenges and systematically validate it in phantom and human in-vivo experiments. We demonstrate that the improvements enable the reliable detection of magnetic field changes caused by the TES current flow in the human brain at a sensitivity level of 84 pT for a resolution of 2×2×3 mm^3^ and a 4:20 min measurement duration.

## METHODS

### Subjects

We have recruited 8 healthy volunteers and performed four successive experiments, as described in detail below. One of the volunteers participated in each experiments, one in experiments 1-3, one in only experiment 1, and one in experiments 2 and 3. The remaining four volunteers participated only in the last experiment 4. The participants had no previous histories of neurological or psychiatric disorders and were screened for contraindications to MRI and TES. Written informed consent was obtained from all participants prior to the scans. The study complied with the Helsinki declaration on human experimentation and was approved by the Ethics Committee of the Medical Faculty of the University of Tübingen, Germany and the Capital Region of Denmark.

### Measuring current-induced magnetic fields by gradient-echo MRI

All experiments were performed in 3T MR scanners (MAGNETOM Prisma, SIEMENS Healthcare, Germany) equipped with 64-channel head coils. The MR signals from each channel were combined with an adaptive-combine algorithm (34). We employed a gradient-echo-based steady-state MRI pulse sequence (11) with constant tip-angle RF excitation pulses repeating at a constant time interval *T*_*R*_ and synchronized TES currents to create two steady-state magnetization states that are modulated by positive (+) and negative (−) current-induced magnetic fields (Fig. 2a). We used the steady-state MR signals acquired in each of the *n*^th^ echo intervals to reconstruct a phase difference image ∠*M*_*n*_^+^ − ∠*M*_*n*_^−^ = 2*γ*Δ*B*_*z,c*_*T*_*E*_ (*n*) + Δ*φ*_*ss*_, where *γ* is the proton’s gyromagnetic ratio, *T*_*E*_(*n*) the echo time, and Δ*φ*_*ss*_ the steady-state phase difference just after the excitation. Exploiting the linear dependence of Δ*φ*_*ss*_ on Δ*B_z,c_* for tiny TES currents, we derived the corresponding image of the current-induced magnetic field change Δ*B_z,c_*^*n*^ = (∠*M*_*n*_^+^ − ∠*M*_*n*_^−^)/*m*_*n*_, where the slope *m*_*n*_ highly depends on the MR sequence and relaxation parameters (echo time *T*_*E*_(*n*), repetition time *T*_*R*_, tip-angle *α*, the spoiling scheme; *T*_1_, *T*_2_, and 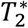) and can numerically be calculated by integrating Bloch equations (11,35,36).

**Figure 2.**
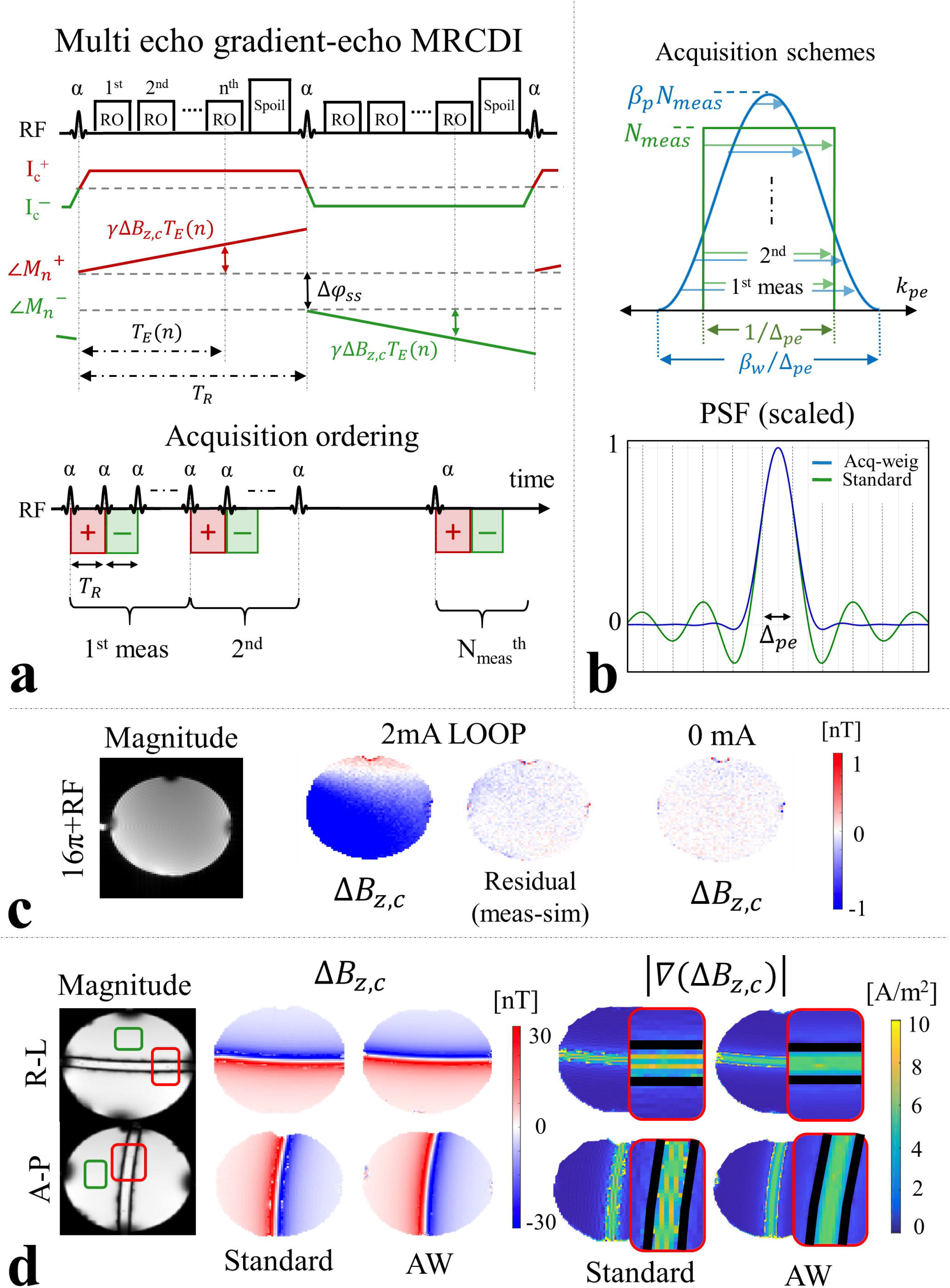
(a) Schematic diagram of a gradient-echo-based MRCDI sequence (see the “Methods” section and (10,11) for details). (b) Unlike standard acquisition, the center of the k-space is measured more frequently than outer k-space for acquisition-weighted MRI. The weights are matched with the post-hoc filter to resolve the ringing in the MR images by suppressing the side lobes in the PSF while preserving the spatial resolution Δ_*pe*_ and SNR. Phantom experiments. (c) Images of Δ*B_z,c_* and its noise floor in a phantom acquired with the combination of RF and 16π gradient spoiling. The residual Δ*B_z,c_* images (measurements for 2 mA current injection with simulations of cable-induced fields subtracted) are very similar to the noise floor images acquired for 0 mA, demonstrating the efficiency of the employed spoiling. For details, see experiment S.1 in the Supplementary Material. (d) Combined MR magnitude images for standard and acquisition-weighted (AW) acquisitions for a phantom with a spherical tube (see experiment S.2 in the Supplementary Material for detail). The cylindrical tube is aligned in right-left (R-L) or anterior-posterior (A-P) direction. The impact of a better PSF is barely visible in the MR magnitude images (the green rectangles show the regions used for SNR calculations reported in Supplementary Table S2; the red rectangles show the positions of the zoomed regions). AW improves the image quality and the noise floors of the Δ*B_z,c_* image, e.g. spurious bright spots in the Δ*B_z,c_* image close to the tubes are resolved. The improvements in spatial resolution are more obvious in images of the norm of the gradient of the current-induced field images |∇(Δ*B_z,c_*)|: The spatial derivative operation relevant for MRCDI reconstruction amplifies signal variations due to ringing for standard acquisition. This is resolved in the results obtained with AW. The signal-void tube is concealed with black rectangles.

The numerical slopes used for the Δ*B_z,c_*^*n*^ measurements differ depending on the two spoiling techniques employed in our MRCDI sequence: First, we employed constant phase RF excitation and systematically changed the spoiler gradient areas ensuring different amount of intra-voxel phase dispersions *φ*_*sp*_=[2π- 32π] to test the impact of Δ*φ*_*ss*_ simulation accuracy on the Δ*B_z,c_* measurements. Here, we estimated the slopes *m*_*n*_ based on constant nominal sequence parameters and approximate brain tissue relaxation parameters at 3T similar to (11). We also incorporated a 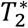 distribution obtained by fitting a perfectly spoiled gradient-echo MRI signal model with a decaying exponential *M*_*ss*_(*t*) = *M*_*ss*_(*t* = 0) ∙ 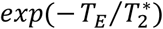 to the acquired MR magnitude images. Second, we combined a strong spoiler gradient with a well-known RF-spoiling method that varies the phase of the *j*^th^ applied RF pulses according to the phase-cycling scheme *φ*_*j*_ = *φ*_*j*−1_ + *jφ*_0_ with *φ*_0_=50° phase increments (37). Here, we used a nulled steady-state phase difference Δ*φ*_*ss*_ = 0 in Δ*B_z,c_*^*n*^ calculations assuming perfect spoiling.

Finally, each of the calculated Δ*B_z,c_*^*n*^ images were systematically weighted and then combined to minimize the noise sensitivity of the combined Δ*B_z,c_* image (10,11,38). The underlying theory can be found in the Supplementary Material. We corrected the combined Δ*B_z,c_* images for the stray magnetic fields that are induced by cable currents similar to our previous study (31). This involved cable tracking using an ultra-short *T*_*E*_ sequence, and subtraction of the corresponding calculated fields.

### Measurement procedures and MRCDI experiments

We employed a transcranial electric stimulation device (DC-STIMULATOR MR, NeuroCare Group GmbH, Germany) in two different setups (Fig. 1). LOOP-SETUP: The generated currents were flowing in a cable loop placed around the head. TES-SETUP: The generated currents were injected into the head via round scalp electrodes for two different montages, anterior-posterior (A-P) and right-left (R-L). We employed 2 mA baseline-to-peak currents for the experiments using LOOP-SETUP and 1 mA for TES-SETUP and imaged two single slices in superior and inferior parts of the brain, respectively. The study comprises four successive experiments measuring the human brain in-vivo:

1. We introduce an optimized spoiling scheme and test how it influences the MRCDI sensitivity and quality.
2. We tested the impact of acquisition weighting on the sensitivity and the resolution of the Δ*B_z,c_* measurements, and the accuracy of the current flow reconstructions.
3. We explored if undesired physiological MR signal fluctuations can be tracked by navigators and used for data quality assessment.
4. We compared our original (10) and improved MRCDI methods. The experiments were performed for two different electrode montages ensuring current injection profiles in anterior-posterior (A-P) and in right-left (R-L) directions.

We used the LOOP-SETUP in the experiments 1-3 and the TES-SETUP in experiment 4 (Fig. 1). Experiments 1 and 2 were preceded by pilot experiments in phantoms, which are described in the Supplementary Material.

### Experiment 1: Importance of proper spoiling in steady-state MRCDI

We used the LOOP-SETUP (Fig. 1) to compare the sensitivity and accuracy of our gradient-echo-based MRCDI method for different spoiling schemes in three subjects. Imaging parameters were FOV = 224×183 mm^2^, *α*=30°, *T*_*E*_ = [5.6, 14.4, 23.2, 32, 40.8, 49.6] ms, *T*_*R*_=80 ms, and an imaging matrix of 112×92. The measurements were repeated *N*_*meas*_ = 16 times to increase the signal-to-noise-ratio (SNR). The experiments were repeated for five different spoiler gradient strengths to ensure intra-voxel phase dispersions of *φ*_*sp*_=[2π, 4π, 8π, 16π, and 32π] without RF spoiling, and the experiment with *φ*_*sp*_=16π was then tested with RF-spoiling.

### Experiment 2: Improving the spatial resolution and sensitivity by acquisition weighting

We used the LOOP-SETUP to compare two different acquisition strategies in three subjects (standard vs. acquisition weighting; Fig. 2b). We explored the influence of the point spread function (PSF) quality on Δ*B_z,c_* measurements and tested the impact of acquisition weighting on the sensitivity and resolution of the MRCDI measurements. The total scan time was kept close to 4:20 mins for both acquisition strategies. Imaging parameters were FOV = 224×183 mm^2^, *α*=30°, *T*_*E*_ = [5.6, 14.4, 23.2, 32, 40.8, 49.6, 58.4, 67.2] ms, *T*_*R*_=80 ms, and combined *φ*_*sp*_=16π and RF spoiling:

- Standard acquisition. We employed a standard k-space acquisition scheme in a finite-rectangular window corresponding to a pre-set nominal resolution Δ_*pe*_=2 mm in the phase encoding direction and kept the total number of repetitions identical *N*_*meas*_ = 16 for each of the phase encoding lines, i.e. uniform weighting (this acquisition scheme results in the standard sinc-shaped PSF that is known to cause ringing artefacts). An imaging matrix of 112×92 was used.
- Acquisition weighting. As a second strategy, we employed acquisition weighting (AW) ensuring an SNR-efficient PSF improvement (32,33). The k-space data was acquired in a ~1.6 times broader window and filtered with a Hanning window (*h*(*k*_*pe*_) = *β*_*p*_(1 + *cos*(2π*k*_*pe*_Δ_*pe*_/*β*_*w*_))/2, where *β*_*w*_ = 1.6 determines the width of the filter, and *β*_*p*_ = 1.2 the peak value) in both phase encoding and readout direction. The number of measurements was modified systematically to match the applied filter in the phase encoding direction, which ensured a near-flat noise power after filtering (Fig. 2b). An imaging matrix of 176×144 was used to match the actual resolutions of both cases.

### Experiment 3: Using navigators for data quality assessment

To enable the continuous tracking of global field fluctuations due to physiological noise and subject motion, we modified the acquisition-weighted sequence of experiment 2 and replaced the first echo within every *T*_*R*_ with a 0-D navigator while keeping the other parameters unchanged. We quantified the resulting SNR decrease by reanalyzing the data acquired in experiment 2, but discarding the first echo. Then, we used the LOOP-SETUP in three subjects to test whether the navigator signal could successfully track intentional jaw movements.

### Experiment 4: Human in-vivo brain MRCDI for two different current injection profiles

We compared the improved MRCDI method against our original method (10) that to our knowledge provides the best current-induced field measurement efficiency in-vivo documented in the literature so far (~0.1 nT in a ~9 mins scan). We performed experiments (Fig. 1; TES-SETUP; 1 mA) in five subjects for two different current injection profiles in the A-P and R-L directions. Our original MRCDI method parameters were: Number of measurement repetitions *N*_*meas*_ = 36 (here reduced to 18 due to total scan time restrictions), *T*_*R*_ = 80 ms, number of gradient echoes *N*_*GE*_ = 5, *T*_*E*_ = [7.5, 19.8, 31.9, 44, 56.2] ms, *α*=30°, spoiler gradient *φ*_*sp*_= 4π, imaging matrix 112×90, voxel size 2×2×3 mm^3^. A chemical-shift-selective (CHESS) fat suppression technique was applied in the experiment (43).

The improved method with acquisition weighting was used as described above for experiment 2. The first echo was replaced with a 0-D navigator. We did not perform fat suppression as multiple gradient-echo acquisition entails a sufficiently high bandwidth, and instead used the period for acquisition of extra echoes. Both experiments with our original and improved method used the same FOV = 224×183 mm^2^ and the total scan times kept at ~4:20 mins for direct comparison of the sensitivity and resolution. In this study, none of the measurements were discarded as no abnormal signal fluctuations were observed in the navigator signals.

### Noise floor measurements in the ΔB_z,c_ images

In each of the experiments, Δ*B_z,c_* control measurements without any current injections were performed and used to calculate the noise floors. For the phantom experiments, a mostly homogenous region-of-interest (ROI) was selected and a Gaussian distribution was fitted to the Δ*B_z,c_* measurements to evaluate the standard deviations analogous to our previous study (10). The accuracy of the used noise floor estimation method was validated for the phantom experiments by means of a direct comparison with theoretically calculated noise floors obtained from MR magnitude SNR (please see Supplementary Figure S3 and Tables S1&S2). The noise floors were calculated in the entire brain tissue masks for the human in-vivo case.

### Simulating the magnetic fields caused by cable-currents

Prior to the MRCDI measurements a 3D high-resolution structural scan based on the Pointwise Encoding Time reduction with Radial Acquisition (PETRA) sequence (44) was performed to delineate the cable paths. The used parameters were: The number of slices *N*_*sli*_=320, imaging matrix 320×320, voxel size 0.9×0.9×0.9 mm^3^, *T*_*R*_ = 3.61 ms, *T*_*E*_ = 0.07 ms, inversion time *T*_*I*_ = 0.5 s, tip-angle *α*= 6^°^, bandwidth BW=359 Hz/pixel, and turbo factor 400.

In the MRCDI experiments performed with LOOP-SETUP, the fully delineated cable path was used to simulate the expected Δ*B_z,c_* fields based on the Biot-Savart law similar to our previous studies (10,31). The simulated Δ*B_z,c_* fields were subtracted from the measurements to obtain the residual Δ*B_z,c_* images that are ideally zero. The residuals were then compared with the control measurements of the noise floor for no current injection. In the MRCDI experiments with TES-SETUP, the delineated cable paths up until the center of the scalp electrodes were used to simulate and correct for the stray magnetic fields due to the cable currents.

### FEM simulations of the current flows and current-induced magnetic fields

For each participant in experiment 4, we performed additional high-resolution *T*_1_- and *T*_2_-weighted structural scans to inform personalized finite element method (FEM) simulations of the TES current flow and the current-induced magnetic fields (see (10) for the MR sequence details). The structural scans were used to create volume conductor models using the headreco pipeline (45) in the open-source software SimNIBS 3 (www.simnibs.org; (46)). The models were composed of five tissue compartments differing in their ohmic conductivities: Gray matter (GM - 0.275 S/m), white matter (WM - 0.126 S/m), cerebrospinal fluid (CSF - 1.654 S/m), skull (0.010 S/m) and scalp (0.465 S/m). The positions of the rubber TES electrodes were determined from the PETRA measurements. The electrodes (29.4 S/m) were modelled as discs with 5 mm thickness, 5 cm diameter. The thickness of the gel layers (0.37 S/m) between the electrodes and the scalp were determined from the PETRA images. Dirichlet boundary conditions for the electrostatic potential were applied at the electrode surfaces (49). The current flow distributions for A-P and R-L current injection profiles were simulated for 1 mA current injections, and the current flow simulations were then used to calculate the current-induced magnetic field Δ*B_z,c_* distributions based on Biot-Savart simulations (50).

### Current flow reconstructions

A uniform conductivity of 1 S/m was assigned to the electrode pads, gel layer and all compartments of the head models for the subjects in experiment 4 to obtain 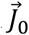 and Δ*B_z,c_*^0^. The projected current densities 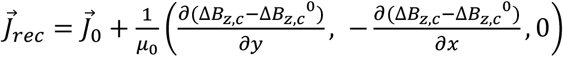 were then reconstructed for the Δ*B_z,c_* measurements and also for the Δ*B_z,c_* simulations with literature conductivity values (51,52). We used a median filter of 3×3 to eliminate high frequency noise in the Δ*B_z,c_* measurements, and the central difference approximation for the directional derivative operations in the reconstruction algorithm. The current flow reconstructions based on the measurements with our original and improved method were compared in terms of quality.

As the reconstruction approach employs spatial gradient operations on the measured Δ*B_z,c_* image, we visualized the norm of its gradient |∇(Δ*B_z,c_*)| as additional quality index when comparing different measurements taken with the LOOP-SETUP.

## RESULTS

### Subject sensations

In the experiments with LOOP-SETUP, none of the subjects reported any side effects or any discomfort. In the experiment 4 employing TES, all subjects reported phosphenes and a slight tingling sensation near the electrodes that disappeared after a short while for both electrode montages A-P and R-L. None of the subjects reported any further discomfort due to TES.

### Experiment 1: Importance of proper spoiling in steady-state MRCDI

The spoiling scheme was initially optimized using tests in phantoms (experiment S.1 in the Supplementary Material). The combination of RF spoiling and gradient-spoiling with *φ*_*sp*_=16π demonstrated the lowest noise floors. Specifically, the residual noise floor after subtracting the fields of the LOOP cables was in the same range as for control measurements without current injection (Fig. 2c). This spoiling scheme was therefore used for the further testing in humans described below. For the control measurements, the estimated noise floors agreed well with theoretically calculated noise floors obtained from MR magnitude SNR (Table S1), validating the employed method for noise floor estimations.

The results for the original and new spoiling schemes are exemplarily shown in Figure 3a for the first subject. No severe artifacts are observed in any of the MR magnitude images. However, gradient spoiling with *φ*_*sp*_=4π results in erroneous Δ*B_z,c_* fields in CSF regions, likely due to steady-state modelling errors. These errors are resolved by the optimized spoiling (16π & RF). The standard deviations of the residual Δ*B_z,c_* images are listed in Table 1a to quantify the noise floors in all 3 subjects. In case of optimal spoiling (16π & RF), the standard deviations were only slightly increased by 7% when compared against the control measurements without any currents. In summary, correcting the cable-induced stray fields works very well when proper spoiling is applied and hardly impacts the noise floor in the corrected images. The complete results for all subjects and spoiling schemes can be found in the Supplementary Figure S4.

**Figure 3.**
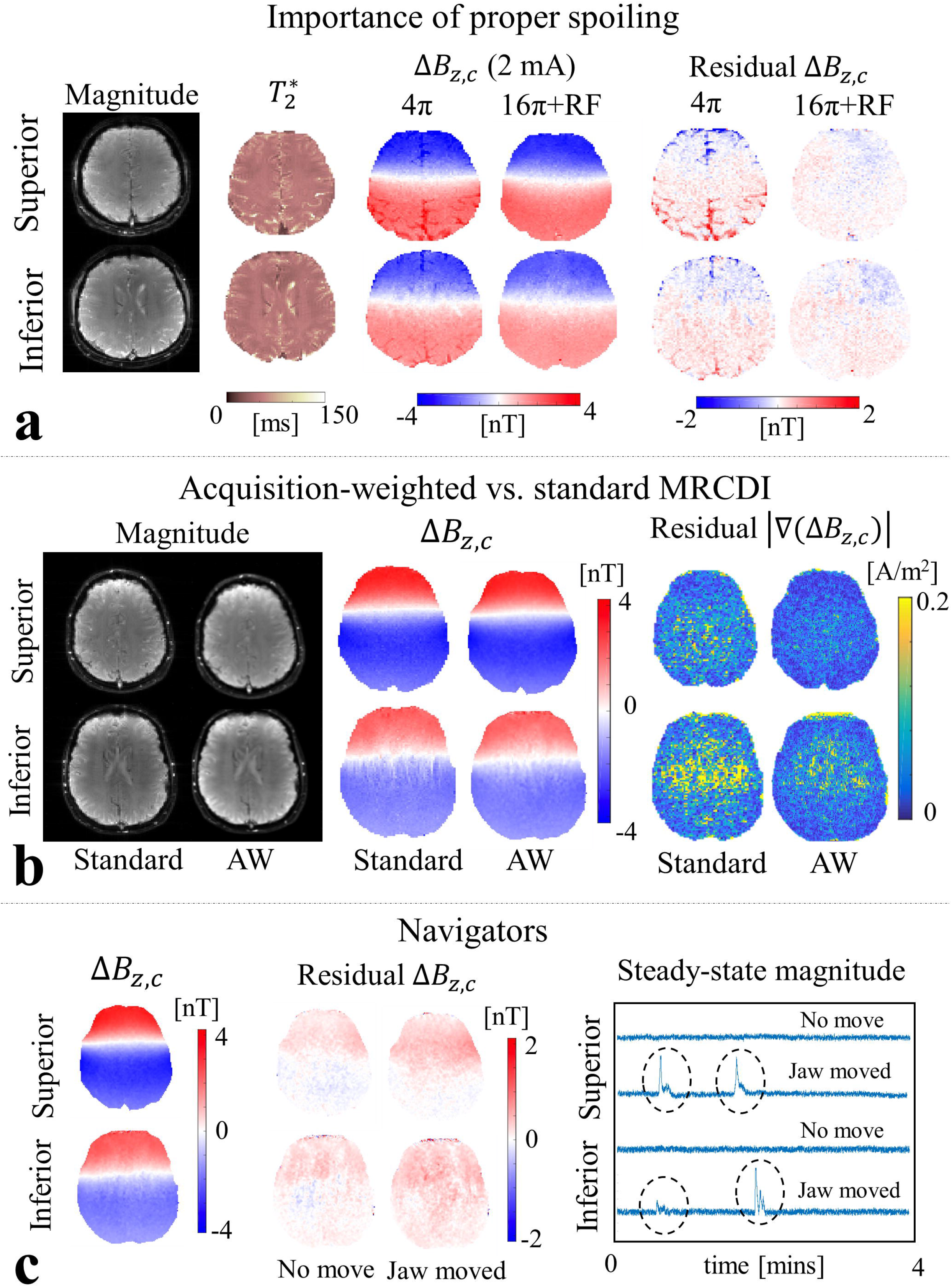
Results for the first subjects in experiments 1 to 3. (a) Experiment 1: MR magnitude and 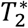 images exemplary shown for the combination of RF and *φ*_*sp*_=16π spoiling. 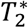 is found to be significantly higher in CSF regions compared to brain tissue. Δ*B_z,c_* measurements and residual images (ideally zero) clearly depict erroneous Δ*B_z,c_* fields in CSF regions for the *φ*_*sp*_=4π spoiling of our original method, likely due to steady-state modelling errors. Combined RF and *φ*_*sp*_=16π spoiling resolves this problem. (b) Experiment 2: Employing AW improves the quality of the MR magnitude and Δ*B_z,c_* images compared to the standard acquisition scheme (ringing-caused fluctuations are suppressed by means of AW). Images of the norm of the gradient |∇(Δ*B_z,c_*)| obtained from Δ*B_z,c_* residuals (ideally zero) more clearly demonstrate the improvement in the noise floors for the AW measurements. (c) Experiment 3: Navigator measurements when the subject performs no intentional movement (No move) or intentional jaw movement twice during the scan (Jaw moved). No obvious quality changes are visible in the Δ*B_z,c_* measurements, but the noise floors increase with jaw movement (Table 1c). The MR magnitude signal acquired from the navigator clearly fluctuates during jaw movements, and otherwise stays constant (note that the first 8 secs are not plotted as the MR magnitude enters steady-state within this period).

**Table 1.**
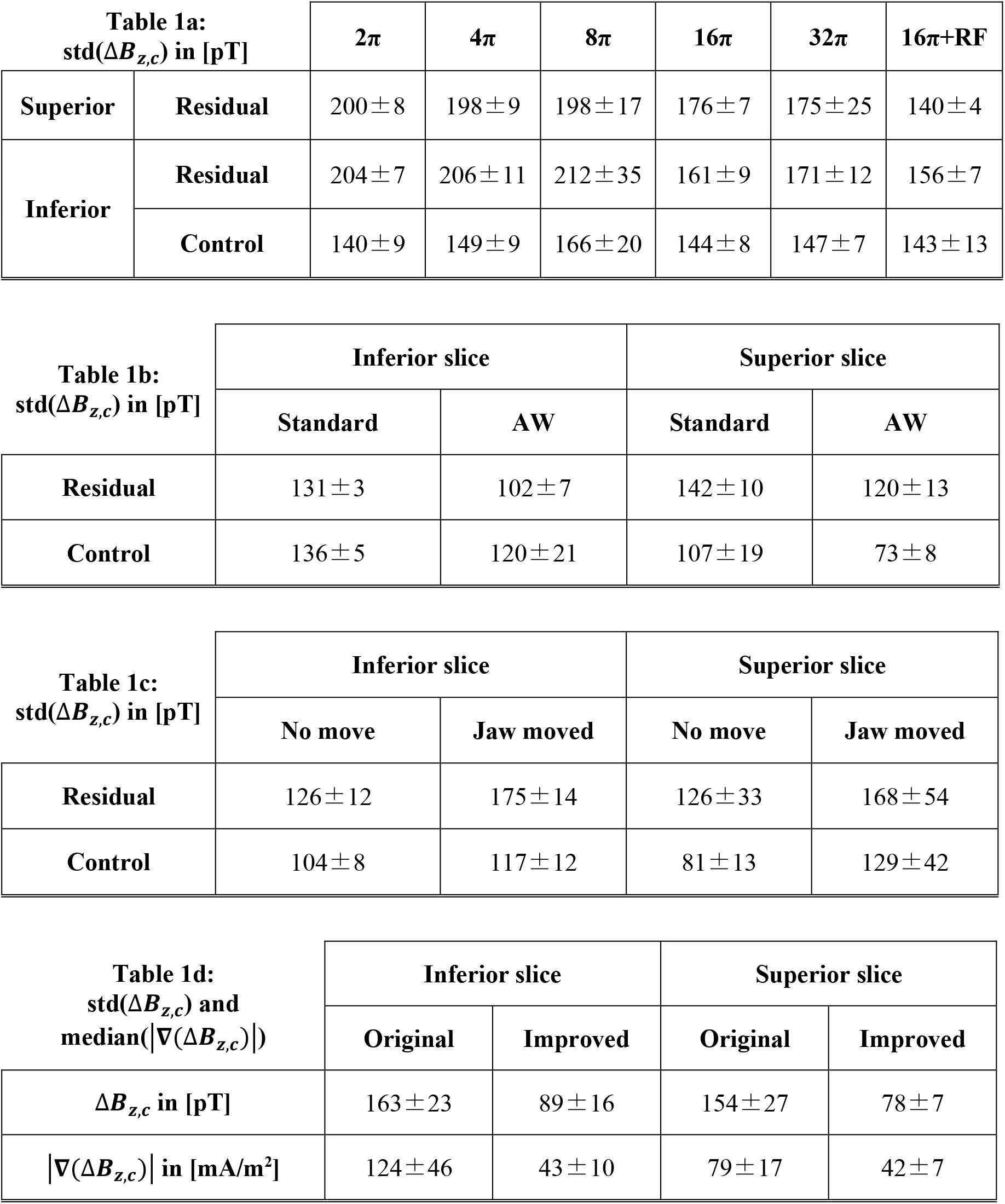
The average standard deviations of Δ*B_z,c_* images and their standard deviations across three subjects are given in [pT]. The Δ*B_z,c_* simulation based on the reconstructed cable paths are subtracted from the Δ*B_z,c_* measurements to obtain residual images. (a) Experiment 1: To explore the importance of proper spoiling in the Δ*B_z,c_* measurements, the standard deviations of the Δ*B_z,c_* residual images and of images from control measurements with no currents are compared for different spoiling schemes ([2π-32π] intra-voxel dephasing employed without RF-spoiling and 16π dephasing combined with RF-spoiling). The combination of RF and 16π gradient spoiling performs best among the tested schemes in terms of standard deviation. (b) Experiment 2: Comparison of standard acquisition with no post-filtering applied and acquisition weighting (AW). Employing AW in the phase-encoding direction significantly improves the Δ*B_z,c_* measurement sensitivity. (c) Experiment 3: Impact of an intentional subject movement (jaw movement twice during the scan) on the Δ*B_z,c_* measurement sensitivity. Significant increases of the noise floor are observed in both residual and control Δ*B_z,c_* measurements due to the subject movement. (d) Experiment 4: Our original method used in (10) is directly compared with our improved method in terms of Δ*B_z,c_* measurement sensitivity and median of the norm of the current-induced field gradients |∇(Δ*B_z,c_*)|. The average standard deviations of Δ*B_z,c_* images and their standard deviations across five subjects are given in [pT]. The average median values of |∇(Δ*B_z,c_*)| images and their standard deviations across five subjects are given in [mA/m^2^]. Our improved method demonstrates more than 2-fold sensitivity increase for a matched total scan time.

### Experiment 2: Improving the spatial resolution and sensitivity by acquisition weighting

Acquisition weighting was initially optimized by tests in phantoms (experiment S.2 in the Supplementary Material) and demonstrated to decrease the noise floors of the Δ*B_z,c_* images. It also resolved ringing artifacts caused by a poor PSF of the standard acquisition (Fig. 2b and d). For the tests in humans, the results of the first subject are exemplarily shown in Figure 3b (see Supplementary Figure S5 for complete results of all subjects). No severe artifacts are observed in the MR magnitude images. The current-induced magnetic field images qualitatively improve and the spurious field fluctuation near ventricles along the phase encoding direction vanishes in the acquisition-weighted images.

Acquisition weighting consistently reduces the noise floors of Δ*B_z,c_* and of the norm of its gradient |∇(Δ*B_z,c_*)| for all subjects, in both the residual images and control measurements without currents. On average among the three subjects, the noise standard deviations are reduced by 28% for the inferior slice and 18% for the superior in the residual Δ*B_z,c_* images; and by 13% for the inferior slice and 46% for the superior in the control measurements without currents (Table 1b).

### Experiment 3: Using navigators for data quality assessment and exploring the influence on sensitivity

Discarding the first echo in the combined Δ*B_z,c_* measurements of Experiment 2 causes less than 1% increase in the noise floor, which demonstrates that the first echo can be replaced with a navigator without penalty on SNR.

The Δ*B_z,c_* measurements of the first subject in experiments with and without intentional jaw movements and the influence of the movements on the steady-state navigator signal magnitude are shown in Figure 3c (see Supplementary Figure S6 for the results of all subjects). The movements consistently cause peaks in the magnitude signal in all experiments (indicated by black dashed circles), while there are no significant phase variations observed in the navigator signal.

Jaw movements do not have a visually observable effect on the quality of the current-induced magnetic field Δ*B_z,c_* and MR magnitude images. However, on average among the three subjects (Table 1c), it increases the standard deviations by 39% for the inferior slice and 34% for the superior slice in the residual Δ*B_z,c_* images; and by 13% for the inferior slice and 60% for superior slice in the control measurements without current injection.

### Experiment 4: Human in-vivo brain MRCDI for two different current injection profiles

Figure 4a (subject 1) and 5 (subject 2-5) compares the Δ*B_z,c_* measurements acquired with our original method (10) against our improved method. No significant artefacts are observed in the acquired MR magnitude images. Our improved method lowers the Δ*B_z,c_* noise floors in the 0 mA results consistently for every subject. In particular, it resolves the severe artifacts near ventricle regions in the inferior slice that occur for our original method (please note that only superior slices were measured in our initial study (10)). On average across the five subjects, the improved method exhibits an almost two-fold sensitivity increase over our original method, with noise standard deviations in the Δ*B_z,c_* images being as low as 78 pT in the superior slices and 89 pT in the inferior slices (Table 1d). The better quality of the improved method translates to lower noise floors in the images of the norm of the gradient |∇(Δ*B_z,c_*)| (0 mA results in Figure 4b and 6). Particularly, severe noise near the ventricles in the inferior |∇(Δ*B_z,c_*)| images is consistently avoided. On average across the five subjects, the improved method increases sensitivity 2.4-fold in the |∇(Δ*B_z,c_*)| images, with noise medians being as low as ~43 mA/m^2^ in both slices (Table 1d).

**Figure 4.**
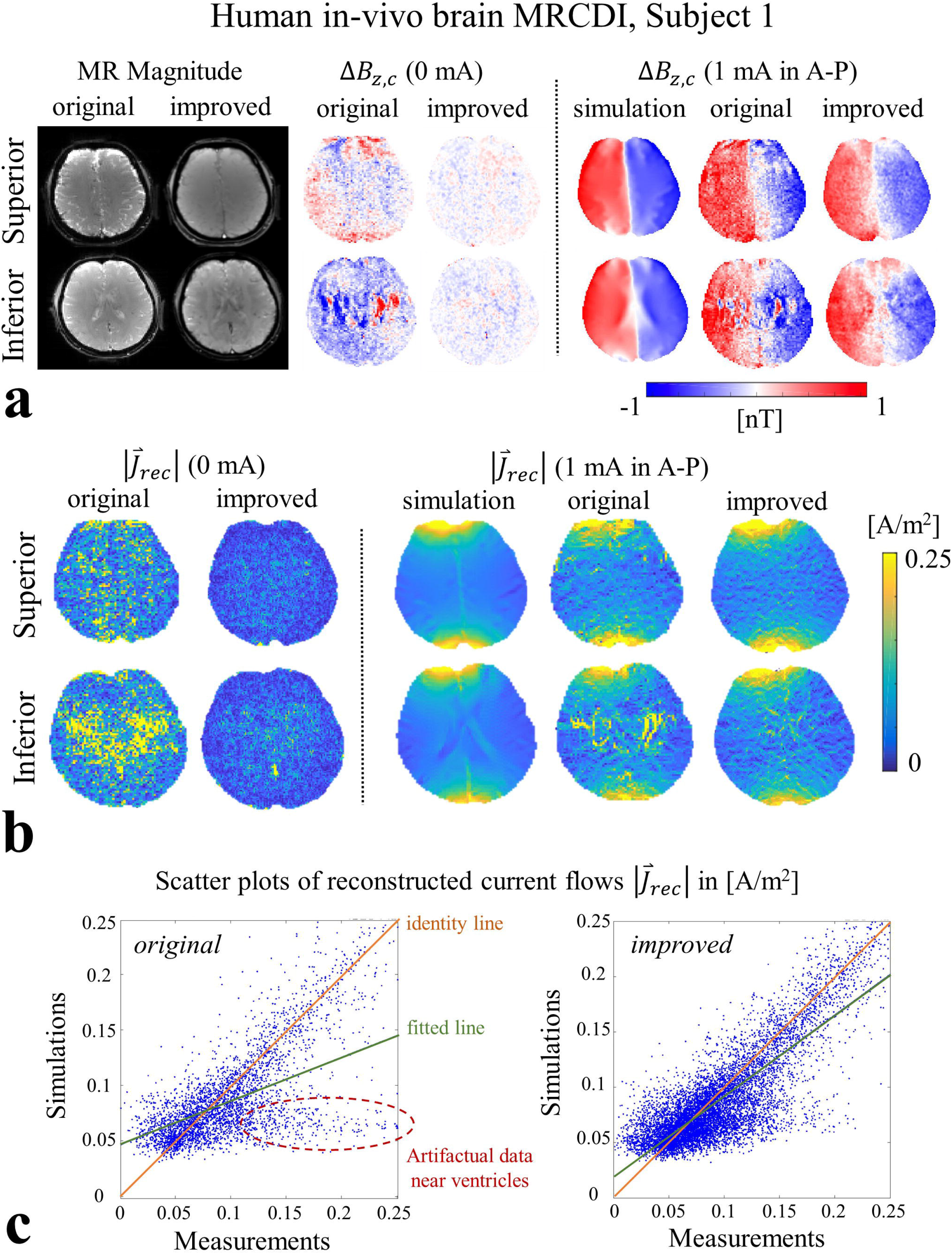
Experiment 4: Results for the first subject for the direct comparison of the final version of our improved MRCDI measurement method with our original method (10). (a) The improved method significantly reduces the Δ*B_z,c_* noise floors (0 mA results) and enhances the image quality and resolution of the Δ*B_z,c_* measurements (1 mA A-P current injection). It also successfully resolves spurious Δ*B_z,c_* variations occurring near ventricles. The Δ*B_z,c_* simulations and measurements show similar distributions, and are in the same range. (b) SNR and image quality changes are more clearly observed in the current flow reconstructions based on Δ*B_z,c_* measurements and simulations, where noise floors are strongly reduced for our improved method. The reconstructed current flows from the simulations are similar to the measurements. (c) The scatter plots of the inferior slice results clearly demonstrate the relevance of the improved measurement method. The fitted line (green) approaches the identity line (orange) that is the ideal case. Our improved method also resolves the cluster (red) corresponding to the artifacts near ventricles.

Also for the measurements with 1 mA TES current injection, the improved method consistently achieves a higher SNR and better image quality of the results. This is apparent in both the Δ*B_z,c_* images (A-P montage in Figure 4a and 5; R-L montage in Supplementary Figure S7) and the current flow reconstructions 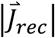 (A-P montage in Figure 4b and 6; R-L montage in Supplementary Figure S8). The improvements are most clearly seen in the regions near the ventricles that suffer from severe artefacts in the results obtained with our original method.

**Figure 5.**
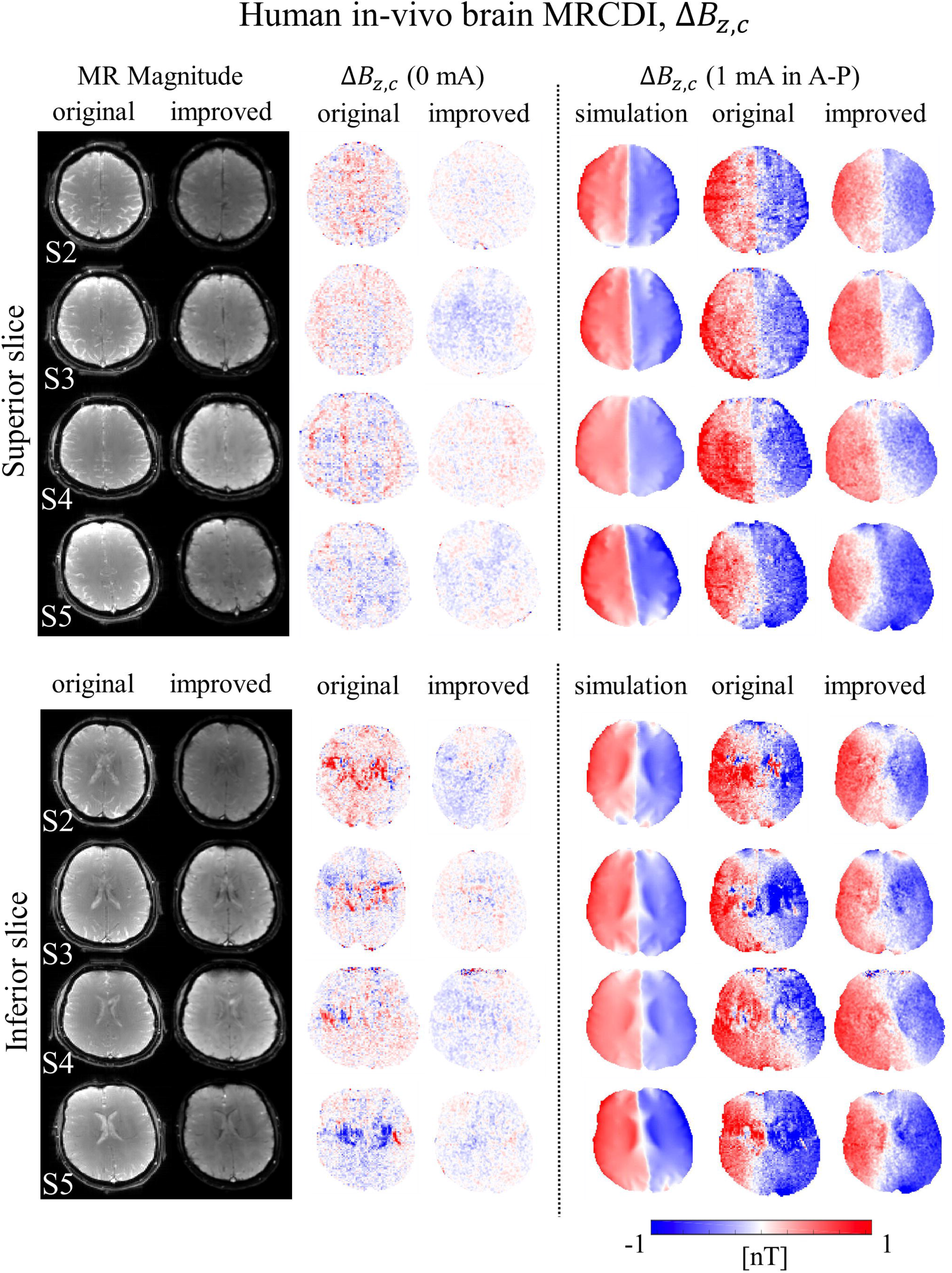
Results for the remaining four subject of experiment 4 for 0 mA and 1 mA A-P current injections (see Figure 4a for the first subject). The improved method significantly reduces the Δ*B_z,c_* noise floors, enhances the image quality and resolution and successfully resolves spurious Δ*B_z,c_* variations near ventricles. The Δ*B_z,c_* measurements and simulations generally agree well, but show slight differences near electrodes. Both simulations and measurements are in the same range. The results are consistent among all five subjects.

**Figure 6.**
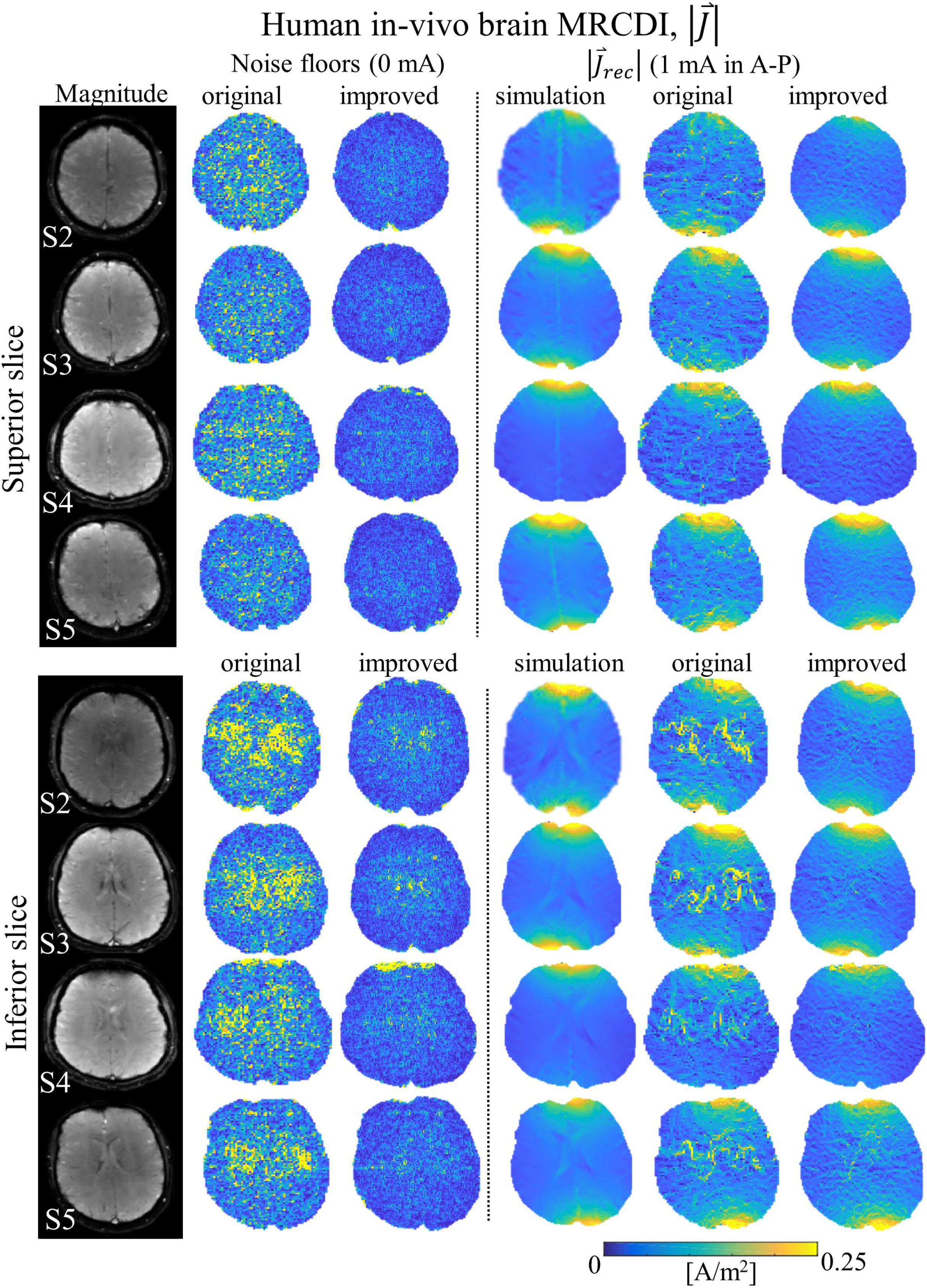
Current flow reconstructions for the remaining four subject of experiment 4 for 0 mA and 1 mA A-P current injections (see Figure 4b for the first subject). Noise floors decrease markedly for the improved method and the artefacts near ventricles are resolved. This consistently improves the correspondence between the current flows reconstructed from the simulations and the measurements.

The simulations of Δ*B_z,c_* based on realistic individualized head models show very similar spatial distributions to the measurements with the A-P electrode montage, whereas clear differences are observed for some cases with the R-L montage (1 mA results in Figures 4a, 5 and S7). For both of the electrode montages, the simulations deviate from the measurements near electrode regions. We tested quantitatively how well the simulations fit to the measurements by linearly regressing the simulated against the measured Δ*B_z,c_* values, analogous to our previous study (10). The intercepts *β*_0_ are in acceptable ranges and the slopes *β*_1_ are closer to one for our improved method (Table 2a). Our improved method demonstrates consistent increases in the coefficients of determination for all subjects (superior slice: Δ*R*^2^=0.07 for A-P and Δ*R*^2^=0.08 for R-L; inferior slice: Δ*R*^2^=0.18 for A-P and Δ*R*^2^=0.14 for R-L, averaged across subjects). The regression analysis was repeated for the current flow images reconstructed from the measured and simulated Δ*B_z,c_* results. The intercepts *β*_0_ are in acceptable ranges and the slopes *β*_1_ are getting closer to one for our improved method (Fig. 4c and Table 2b). It again shows a consistent increase in the coefficients of determination (superior slice: Δ*R*^2^=0.17 for A-P and Δ*R*^2^=0.19 for R-L; inferior slice: Δ*R*^2^=0.32 for A-P and Δ*R*^2^=0.21 for R-L, averaged across subjects).

**Table 2.** Experiment 4: The regression analyses to explore the correspondence between measurements and simulations based on realistic head models are given as averages across the five subjects: (a) Δ*B_z,c_* recordings, and (b) current flow reconstructions 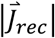. Linear fits of the measurements and simulations for the current injections in A-P and R-L directions are compared for our original and improved methods. The table lists the intercepts *β*_0_, the slopes *β*_1_, and the coefficient of determination *R*^2^ of the fitted linear regression models (± the standard deviations across the five subjects). The significances of the regression models were confirmed using F-tests, with the results being highly significant (p<10^−6^) in all cases. Interestingly, the correspondence to the simulations consistently increases for our improved method, indicating the impact of an improved measurement SNR on the fitting accuracy. In particular, the artefacts observed near ventricles dramatically reduces the correspondence for the inferior slice measurements with our original method.

| Table 2a |  | Original |  |  | Improved |  |  |
| --- | --- | --- | --- | --- | --- | --- | --- |
| | | $\beta_0$<br>in [pT] | $\beta_1$ | $R^2$ | $\beta_0$<br>in [pT] | $\beta_1$ | $R^2$ |
| A-P | Superior | $40 \pm 81$ | $0.83 \pm 0.12$ | $0.77 \pm 0.02$ | $-33 \pm 137$ | $1.12 \pm 0.10$ | $0.84 \pm 0.04$ |
| | Inferior | $28 \pm 74$ | $0.66 \pm 0.12$ | $0.61 \pm 0.06$ | $-92 \pm 74$ | $0.93 \pm 0.11$ | $0.79 \pm 0.03$ |
| R-L | Superior | $2.4 \pm 97$ | $0.41 \pm 0.13$ | $0.31 \pm 0.26$ | $10 \pm 140$ | $0.75 \pm 0.13$ | $0.39 \pm 0.22$ |
| | Inferior | $-31 \pm 105$ | $0.51 \pm 0.09$ | $0.37 \pm 0.05$ | $-61 \pm 120$ | $0.83 \pm 0.19$ | $0.51 \pm 0.09$ |

**Table 2.** Experiment 4: The regression analyses to explore the correspondence between measurements and simulations based on realistic head models are given as averages across the five subjects: (a) $\Delta B_{z,c}$ recordings, and (b) current flow reconstructions $|\vec{j}_{rec}|$ .
| Table 2b |  | Original |  |  | Improved |  |  |
| --- | --- | --- | --- | --- | --- | --- | --- |
| | | $\beta_0$ in<br>[mA/m<sup>2</sup>] | $\beta_1$ | $R^2$ | $\beta_0$ in<br>[mA/m<sup>2</sup>] | $\beta_1$ | $R^2$ |
| A-P | Superior | $22.6 \pm 6.4$ | $0.69 \pm 0.09$ | $0.61 \pm 0.09$ | $13.1 \pm 1.6$ | $0.83 \pm 0.04$ | $0.78 \pm 0.05$ |
| | Inferior | $39.4 \pm 6.7$ | $0.44 \pm 0.06$ | $0.37 \pm 0.07$ | $18.8 \pm 5.2$ | $0.72 \pm 0.10$ | $0.69 \pm 0.06$ |
| R-L | Superior | $19.7 \pm 4.3$ | $0.67 \pm 0.07$ | $0.64 \pm 0.07$ | $11.7 \pm 1.8$ | $0.82 \pm 0.03$ | $0.83 \pm 0.04$ |
| | Inferior | $27.3 \pm 7.2$ | $0.53 \pm 0.1$ | $0.52 \pm 0.13$ | $14.8 \pm 2.1$ | $0.74 \pm 0.04$ | $0.73 \pm 0.06$ |

## DISCUSSION

We propose a new gradient-echo-based MRCDI method with an optimized spoiling scheme, acquisition weighting and navigators to reliably detect the tiny magnetic fields induced by TES in the human brain. It demonstrates a good image quality of the current-induced magnetic field changes, robustness to physiological noise and a sensitivity of 84 pT for a resolution of 2×2×3 mm^3^ in a ~4:20 mins total scan time. The improvements successfully resolved several limitations of our prior approach (10):

First, our experiments in phantoms and humans demonstrated the importance of using stronger spoiling than normally needed for imaging. We used 2 mA currents (baseline-to-peak) in the LOOP-SETUP experiments to unambiguously assess the efficiency of the different spoiling approaches. In contrast, we applied only 1 mA when imaging the current flow induced by TES in the brain and the use of fat suppression was not allowing acquiring later echoes in our previous study (10), so that the impact of insufficient spoiling was far less obvious. However, our new findings suggest that the current-induced field changes around small CSF-filled sulci reported in (10) may be exaggerated. Specifically, the results in (10) might have been influenced by residual effects of CSF flow and by the choice of the relaxation parameters in the steady-state models that is not accurate for CSF regions.

Second, our results demonstrated that acquisition of more echoes instead of employing fat suppression can improve the sensitivity of MRCDI measurements without compromising the image quality. The high bandwidth of the acquisition prevents that signal originating from the scalp and the fatty spongy bone of the skull is shifted into the brain.

Third, acquisition weighting efficiently resolved ringing artifacts caused by the sinc-shaped PSF of standard acquisition schemes. In direct comparison to our original method (10), the new method achieves better sensitivity levels of the Δ*B_z,c_* images and the reconstructed current flow images in half of the scan time. The improved PSF shape seems particularly beneficial for the reconstruction of the current flow images, as this relies on spatial differentiation.

Lastly, our results demonstrated the feasibility of detecting undesired motion-induced signal fluctuations for quality control by replacing the first echo with a navigator, which causes a negligible SNR compromise. Here, it helped us to rule out the possibility that subject motion caused the discrepancies between Δ*B_z,c_* simulations and measurements, which we observed for the R-L electrode configuration in some of the subjects.

### Comparison of measured and simulated fields

The correspondence between measurements and simulations notably increased for our improved compared to the original method, highlighting the relevance of the enhanced image quality.

Interestingly, the correspondence for the current-induced magnetic fields varied between the two electrode configurations (A-P: *β*_1_=1.03, *R*^2^=0.82; R-L: *β*_1_=0.79, *R*^2^=0.45, averaged across the five subjects and the two slice positions). Considering our careful validations of the MR measurement approach, it seems unlikely that this differences arose from limitations in the measured data. Rather, the results indicate that the employed volume conductor models (including the head, electrode pads, and gel) might be less accurate for simulations of the tested R-L electrode montage. Along similar lines, it is worth noting that the measured and simulated magnetic fields deviate clearly from each other near electrode regions in some of the experiments. While beyond the scope of this study, these findings indicate that MRCDI provides useful data to potentially improve the accuracy of the field simulations, e.g. by testing head models with improved anatomical detail (53).

In contrast to the current-induced magnetic fields, the current density images reconstructed from measurements and simulations demonstrated a good correspondence that was similar for both current injection profiles. However, the images generally lacked fine spatially details, which was also caused by the employed reconstruction method, as already observed in our prior study (e.g., seen in Figure 8 in (10)). For this reason, we did not directly use the current densities obtained by the FEM simulations for the comparison, but reconstructed the current densities again from the simulated current-induced magnetic fields to account for the effect of the reconstruction method. We hypothesize that this might impact the correspondence between measurements and simulations. As of now, we thus consider the similarity between measured and simulated current-induced magnetic fields a more reliable measure of the fit of the simulated current flow.

### Prior studies, status, and future work

So far, only few studies have reported successful MR recordings of the magnetic fields of TES currents induced in the human brain (10,54,55), based on MR methods with sufficient phase sensitivity. While a mismatch between simulated and measured current-induced magnetic fields was still apparent as differences in spatial patterns and peak strengths in the results of two these studies (54,55), this could be mostly resolved in (10) by correcting for the stray magnetic fields due to the cable currents. Compared to the results in (10), we here improved the sensitivity further while we also enhanced the spatial resolution, reduced the sensitivity to CSF flow and minimized the effect of different tissue-specific relaxation times on the estimated current-induced magnetic field changes. Those measures strongly improved the quality of the Δ*B_z,c_* images and also resolved spurious artefacts that had previously remained also after correcting for the cable stray fields. The acquisition time was kept rather short here, which leaves room for further reduction of the noise levels by temporal averaging.

Considering the achieved sensitivity levels and robustness to physiological noise, we are optimistic that our improved method will be useful to characterize individual current flow patterns induced by TES. While further advancing the MR imaging methods will still be relevant, we suggest that improving the reconstruction methods is at least similarly important. For example, the reconstruction method employed here (26,56) resulted in rather coarse current density images even when applied to noiseless simulated current-induced magnetic fields. This is in contrast to its performance reported for phantom measurements, and indicates the need to adapt the approach to the human head anatomy. Alternatively, conductivity reconstructions using the Laplacian ∇^2^(Δ*B_z,c_*) are less affected by stray magnetic fields originating from current flow outside the measured region, which might increase the stability of the reconstruction in particular around electrode regions (27,57). However, the Laplacian causes strong noise amplification, so that further improvements of the MR image quality would likely be required to make conductivity reconstructions from human brain measurements feasible for these methods.

The MR phase images could be further improved in several ways. For example, balanced SSFP can in theory provide an even better sensitivity to the TES-induced field changes and is robust against physiological variations. However, its sensitivity to field inhomogeneities makes its use challenging (58). Double echo planar imaging (59) might help to reduce the sensitivity of the MR data to physiological field fluctuations, but likely has lower sensitivity to the current-induced magnetic fields.

Furthermore, ultra-high field MRI, e.g. at 7 or 9.4T, generally offers higher signal-to-noise ratios (60), but also requires careful optimization of the imaging methods to achieve robust results. Another important goal would be the extension to simultaneous multi-slice or volume acquisition to obtain more complete information of the current-induced magnetic field changes. However, maintaining the sensitivity levels, image quality and robustness to physiological noise within a larger brain volume might be challenging.

## CONCLUSIONS

We have demonstrated MRCDI measurements of the human brain with high sensitivity and high spatial resolution, providing high-quality maps of current-induced magnetic fields with minimized artefacts. Image quality and noise levels were confirmed using control measurements without any current flow and measurements of the magnetic field changes caused by currents flowing in a wire loop around the head. The latter proved important to test for non-linear dependencies of the MR signal phase on the current-related magnetic field changes that, if ignored, can cause systematic artefacts in the current-induced magnetic field images. We suggest that the achieved level of image quality and robustness to physiological noise will be beneficial to make MRCDI a reliable and accurate method for characterizing the individual current flow patterns induced by TES.

## ACKNOWLEDGEMENTS

The financial support of the Lundbeck Foundation (grant R288-2018-236 to CG and R244-2017-196 and R313-2019-622 to AT) and the Max Planck Society and the German Research Foundation (Reinhart Koselleck Project, DFG SCHE 658/12) is gratefully acknowledged. HRS holds a 5-year professorship in precision medicine at the Faculty of Health Sciences and Medicine, University of Copenhagen which is sponsored by the Lundbeck Foundation (Grant Nr. R186-2015-2138).

