## Supplementary material for "Sensitivity and resolution improvement for in-vivo magnetic resonance current density imaging (MRCDI) of the human brain"

### *Phantom experiments*

#### *Experiment S.1: Importance of proper spoiling in steady-state MRCDI*

We employed a spherical phantom of 12 cm in diameter. We cut four holes evenly distributed around the equator and attached four 2 cm long pipes to each of the holes to fill the phantom with a saline solution (1.5 g/L NaCl) doped with 0.1 mM $MnCl_2$ to obtain MR relaxation parameters close to the human brain tissues ($T_1$ = 1.1 s, $T_2$=100 ms). We placed a cable loop around the phantom as in LOOP-SETUP (Fig. 1) for the current transmission and compared the sensitivity and accuracy of our gradient-echo-based MRCDI method for different spoiling schemes. The experiments employed the following sequence settings: Field of view FOV = 224×183 mm$^2$, imaging matrix 112×92, tip-angle $\alpha$=30°, echo times $T_E$ = [5.6, 14.4, 23.2, 32, 40.8, 49.6] ms, repetition time $T_R$=80 ms. The measurements were repeated $N_{meas}$= 16 times to increase the signal-to-noise-ratio (SNR). The experiments were repeated with four different spoiler gradient strengths to provide intra-voxel phase dispersions of $\varphi_{sp}$=[4π, 8π, 16π, and 32π]. The $\Delta B_{z,c}$ measurements were compared for each of the gradient spoiling strengths without RF-spoiling, and the experiment with $\varphi_{sp}$=16π was then tested with RF-spoiling. We also performed control measurements without any currents to obtain $\Delta B_{z,c}$ noise floor images.

The phantom results are shown in Figure S1. MR magnitude images are artifact-free and appear as expected (please note that the used spherical phantom has four cylindrical tubes located at top, bottom, right, and left, and the field inhomogeneities near these tubes cause lower $T_2^*$ and thus local MR signal reductions). $T_2^*$ is around 80 ms in the center of the phantom. Interestingly, increasing the spoiler gradients changes the residual $\Delta B_{z,c}$ images, which are the difference images between the measured and simulated current-induced magnetic fields calculated from the delineated cable paths by using the Biot-Savart law.

Visual inspection suggests that the residuals exhibited two distinct spatial components. The first varies smoothly across the image and has a high (negative) correspondence with the current-induced field's spatial distribution. The second is restricted to the outer rims of the phantom and is strong at positions of low $T_2^*$. The former increases for stronger spoilers, and the latter decreases. Combining spoiler gradients ensuring 16π intra-voxel phase dispersion with RF-spoiling optimally minimizes the residual noise floors to 61 pT for a total scan time $T_{tot}$= 4:20 mins. This is comparable with the noise floor of 57 pT obtained in the control measurements without currents (Supplementary Table S1 in "Noise floor calculations" section).

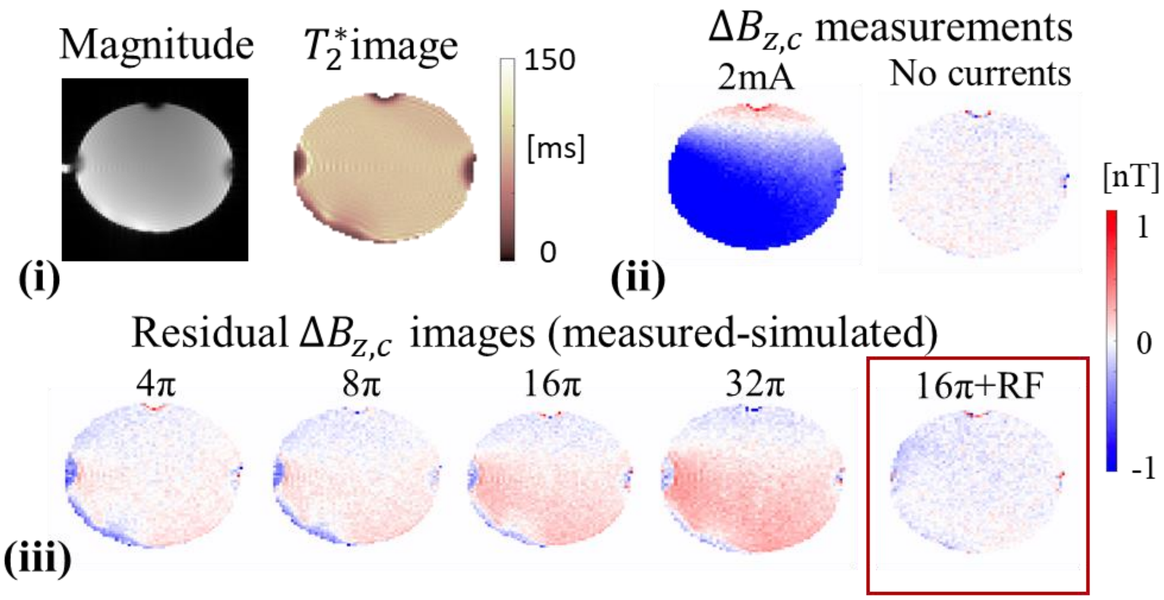


**Supplementary Figure S1.** Experiment S.1: Importance of proper spoiling. (i) Signal losses near pipes are observed in the MR magnitude image of the spherical phantom, mainly due to local field inhomogeneities lowering the $T_2{}^*$. (ii) Images of $\Delta B_{z,c}$ and its noise floor are exemplarily shown for the combination of RF and $16\pi$ gradient spoiling. (iii) Residual $\Delta B_{z,c}$ images (measurements with simulations of cable-induced fields subtracted) are shown for different spoiling schemes, and are expected to be zero ideally. The residual images with no RF-spoiling exhibit strong misestimations of $\Delta B_{z,c}$ in the regions that have low $T_2{}^*$. The combination of RF and $16\pi$ gradient spoiling minimizes the residuals.

*Experiment S.2: Improving the spatial resolution and sensitivity by means of acquisition weighting*

We employed a spherical phantom of 14 cm in diameter. As in the first phantom, we cut four holes evenly distributed around the equator. We attached two 2 cm long pipes to opposite holes to fill the sphere. We inserted a long insulating cylindrical tube (inner diameter=1.1 cm, outer diameter=1.5 cm) directly through the other two holes. The sphere was filled with the same doped saline solution (0.1 mM $MnCl_2$ + 1.5 g/L NaCl) and the cylindrical tube was filled with a conductive gel-like solution (0.1 mM $MnCl_2$ + 1.5 g/L NaCl + 1 g/L TX151). Currents were restrained inside the cylindrical tube only.

The experiments were performed with FOV = 224×183 mm$^2$, tip-angle $\alpha$=30°, echo times $T_E$= [5.6, 14.4, 23.2, 32, 40.8, 49.6, 58.4, 67.2] ms, and repetition time $T_R$=80 ms. The experiments were performed with the optimum spoiling scheme ($\varphi_{sp}$=$16\pi$ combined with RF-spoiling) and repeated for the following three conditions:

a) Standard acquisition. No filtering was applied in the post-processing. The number of measurement repetitions were kept fixed at $N_{meas}$ = 16 for each of the k-space measurements. An imaging matrix of 112×92 was used.

b) Standard acquisition and post-hoc filtering. The k-space data was acquired in a ~1.6 times broader rectangular window and filtered with a Hanning window ($\beta_w$ = 1.6 and $\beta_p$=1.2) in both phase encoding and readout directions. The number of measurements was kept fixed at $N_{meas}$ = 10 for each of the phase encoding lines and an imaging matrix of 176×144 was used to match the total scan time to that of condition (a).

c) Acquisition weighting. The k-space data was acquired in a ~1.6 times broader window and filtered with a Hanning window ($\beta_w$ = 1.6 and $\beta_p$=1.2) as in condition (b). The number of averages was varied systematically to match the applied filter in the phase encoding direction (Fig. 2). An imaging matrix of 176×144 was used.

In each of the conditions the total scan time was kept as close as possible to 4:20 mins.

The experiments were repeated for two different phantom alignments: The cylindrical tube was aligned in the right-left (R-L) and in the anterior-posterior (A-P) directions (superior-inferior tube currents would be parallel to the scanner field and thus induce no measurable current-induced magnetic field component). First, $\Delta B_{z,c}$ noise floors were measured without current injection. Then, currents of 1 mA were injected into the cylindrical tube. As current and conductivity reconstruction algorithms usually apply spatial gradient operations to $\Delta B_{z,c}$, we used the norm of the gradient of the current-induced fields $|\nabla(\Delta B_{z,c})|$ as quality index to characterize how suited the data acquired for the different conditions would be for our application.

Images of $\Delta B_{z,c}$ and the norm of its gradient $|\nabla(\Delta B_{z,c})|$ are shown in Figure 2d. The results illustrate two repeated measurements for two different alignments of the cylindrical tube (R-L and A-P). The quality of MR magnitude images improves in the acquisition-weighted images that have an ameliorated PSF. Ringing affects the $\Delta B_{z,c}$ images in case of current flow through the tube. This is best observed for the images of the norm of the gradient $|\nabla(\Delta B_{z,c})|$, because the derivative operation that is also used by MRCDI and MREIT reconstruction algorithms, emphasizes local signal variations. Specifically, ringing causes two spurious bands at the edges of the tube. Filtering improves the resolution and resolves this issue similar to the AW case.

Applying acquisition weighting with a matched filter provides about 50% decrease in both $\Delta B_{z,c}$ and $|\nabla(\Delta B_{z,c})|$ noise floors on average across measurements for the two alignments. The $\Delta B_{z,c}$ noise floors in a region-of-interest (ROI indicated as green rectangle in Supplementary Figure S2) are given in Supplementary Table S2.

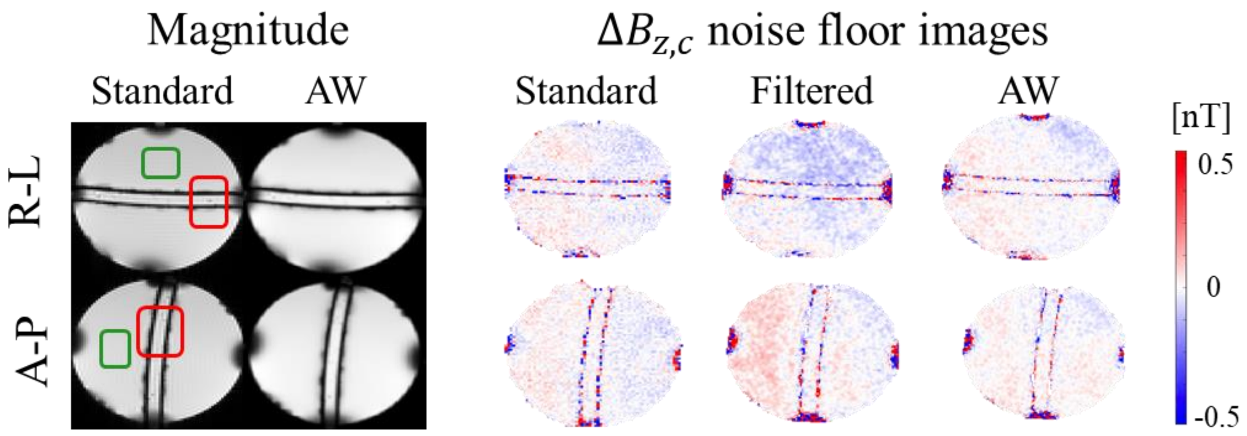


**Supplementary Figure S2.** Experiment S.2: Sensitivity improvement by means of Acquisition weighting (AW). Combined MR magnitude images for standard and acquisition-weighted acquisitions, with alignment of the cylindrical tube in right-left R-L or anterior-posterior A-P direction. The impact of a better PSF is barely visible in the MR magnitude images (the green rectangles show the regions used for SNR calculations reported in Supplementary Table S2; the red rectangles show the positions of the zoomed regions used in Figure 2d). AW improves the noise floors of the $\Delta B_{z,c}$ images.

### *Noise floor calculations*

The standard deviation in a $\Delta B_{z,c}$ measurement based on a single-echo-gradient-echo sequence is given by $std(\Delta B_{z,c}) = 1/\sqrt{2}\gamma T_E SNR$, where $\gamma$ is the gyromagnetic ratio of the proton, $T_{\mathrm{E}}$ the echo time, and $SNR = mean(|M_{ROI}|)/std(|M_{ROI}|)$ the signal-to-noise ratio of the MR magnitude image that can be calculated from the MR magnitude image in a homogenous region-of-interest $|M_{ROI}|$ (23). In this study, we used MR magnitude images for both (+) and (-) currents for a more robust noise floor calculation.

As an example, the MR magnitude and $\Delta B_{z,c}$ images for each of the echoes acquired in Exp S.1 are shown in Supplementary Figure S3. The *SNR* and noise floor values calculated in Exp S.1 are given in Supplementary Table S1, and Exp S.2 are given in Supplementary Table S2.

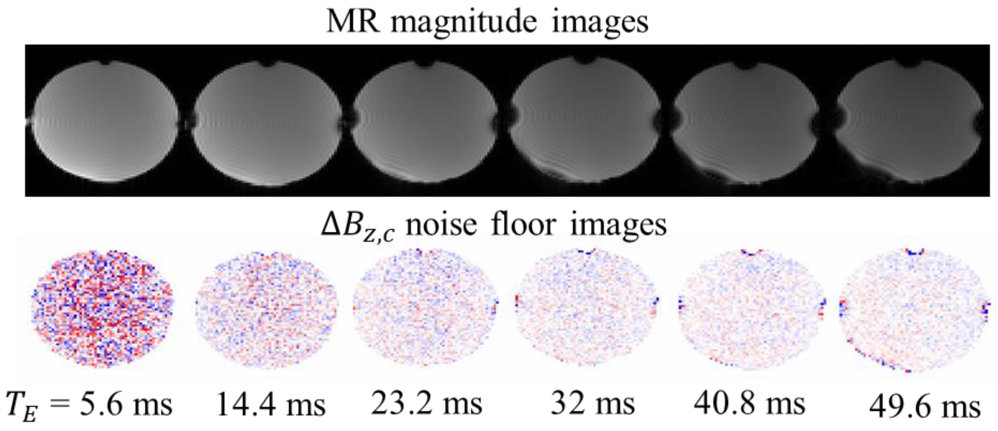


**Supplementary Figure S3.** Experiment S.1: Control experiments without any currents. MR magnitude signal reduces for later echoes and show local signal losses near phantom edges due to the field inhomogeneities as expected. The $\Delta B_{z,c}$ noise images calculated for each of the echoes demonstrate a spatial distribution similar to random noise.

Each of the $\Delta B_{z,c}$ images derived from the $n^{th}$ echo are weighted with its inverse noise variance $1/Var(\Delta B_{z,c}{}^{n}) = 2\,\gamma^2\,T_E(n)^2\,SNR_n^2$ and combined to minimize noise floors (the weights are scaled so that they sum up to one). The noise standard deviation of the combined $\Delta B_{z,c}$ image is $std(\Delta B_{z,c}{}^{comb}) = 1/\sqrt{\sum_{n=1}^{Nechoes} 2\,\gamma^2\,T_E(n)^2\,SNR_n^2}$. In both of the phantom experiments S.1 & S.2, the calculated standard deviations from SNR measurements and the standard deviation measurements based on Gaussian fits strongly agree.

| $T_E$[ms] | 5.6 | 14.4 | 23.2 | 32 | 40.8 | 49.6 | **comb** |
|---|---|---|---|---|---|---|---|
| *SNR* | 1013 | 900 | 795 | 675 | 608 | 624 | **2037** |
| $std(\Delta B_{z,c})$ in [pT] calculated from *SNR* | 467 | 204 | 144 | 123 | 107 | 86 | **52** |
| $std(\Delta B_{z,c})$ in [pT] measurements | 498 | 203 | 134 | 113 | 106 | 98 | **57** |

**Supplementary Table S1**. Experiment S.1: Control measurements without any current. MR magnitude image SNR are given for every individual echo and their weighted combination. The calculated noise standard deviations based on SNR measurements are compared with the noise standard deviations derived from the $\Delta B_{z,c}$ measurements. The theoretical noise floors based on SNR measurements agree well with directly measured $\Delta B_{z,c}$ noise floors.

| | | $T_E$[ms] | 5.6 | 14.4 | 23.2 | 32 | 40.8 | 49.6 | 58.4 | 67.2 | **comb** |
|---|---|---|---|---|---|---|---|---|---|---|---|
| *R-L* | *standard* | *SNR* | 1072 | 928 | 863 | 761 | 641 | 645 | 555 | 488 | **1935** |
| | | $std(\Delta B_{z,c})$ [pT] calculated from *SNR* | 441 | 198 | 132 | 109 | 101 | 83 | 82 | 81 | **37** |
| | | $std(\Delta B_{z,c})$ [pT] measurements | 415 | 198 | 142 | 119 | 99 | 84 | 89 | 80 | **54** |
| | *AW* | *SNR* | 1482 | 1339 | 1217 | 1031 | 965 | 848 | 734 | 587 | **2839** |
| | | $std(\Delta B_{z,c})$ [pT] calculated from *SNR* | 319 | 137 | 94 | 80 | 67 | 63 | 62 | 67 | **28** |
| | | $std(\Delta B_{z,c})$ [pT] measurements | 325 | 141 | 96 | 84 | 71 | 68 | 61 | 59 | **30** |
| *A-P* | *standard* | *SNR* | 1204 | 1114 | 1043 | 888 | 795 | 718 | 685 | 609 | **2398** |
| | | $std(\Delta B_{z,c})$ [pT] calculated from *SNR* | 392 | 165 | 109 | 93 | 82 | 74 | 66 | 65 | **31** |
| | | $std(\Delta B_{z,c})$ [pT] measurements | 394 | 160 | 113 | 99 | 80 | 81 | 73 | 69 | **40** |
| | *AW* | *SNR* | 1523 | 1559 | 1228 | 1018 | 957 | 881 | 836 | 710 | **3126** |
| | | $std(\Delta B_{z,c})$ [pT] calculated from *SNR* | 310 | 118 | 93 | 81 | 68 | 61 | 54 | 55 | **26** |
| | | $std(\Delta B_{z,c})$ [pT] measurements | 289 | 140 | 96 | 76 | 66 | 59 | 56 | 50 | **29** |

**Supplementary Table S2**. Experiment S.2: Anterior-posterior and right-left alignment of the phantom. Control measurements without any current. MR magnitude image SNR are given for every individual echo and their weighted combination. The calculated noise standard deviations based on SNR measurements are compared with the noise standard deviations derived from the $\Delta B_{z,c}$ measurements. The results are given for both acquisition schemes, standard vs. acquisition-weighted.

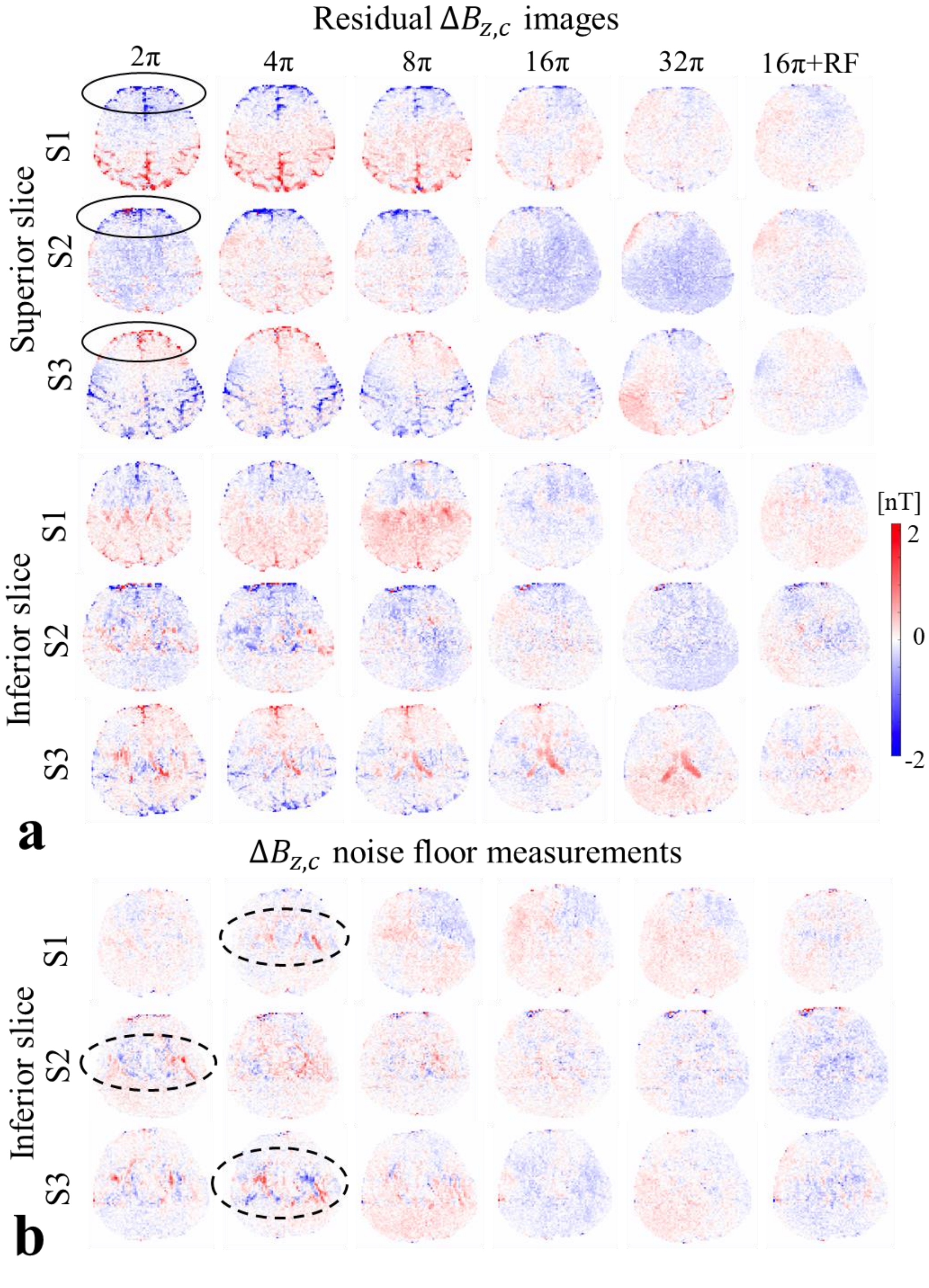


**Supplementary Figure S4.** Experiment 1: Importance of proper spoiling in human in-vivo brain MRCDI. The results of both superior and inferior slice measurements of all subjects. Spurious local increases in the residual $\Delta B_{z,c}$ images near sulci and ventricles are consistently resolved by the combination of RF and $16\pi$ gradient spoiling (simulations subtracted from the magnetic fields measurements performed with LOOP-SETUP; ideally zero).

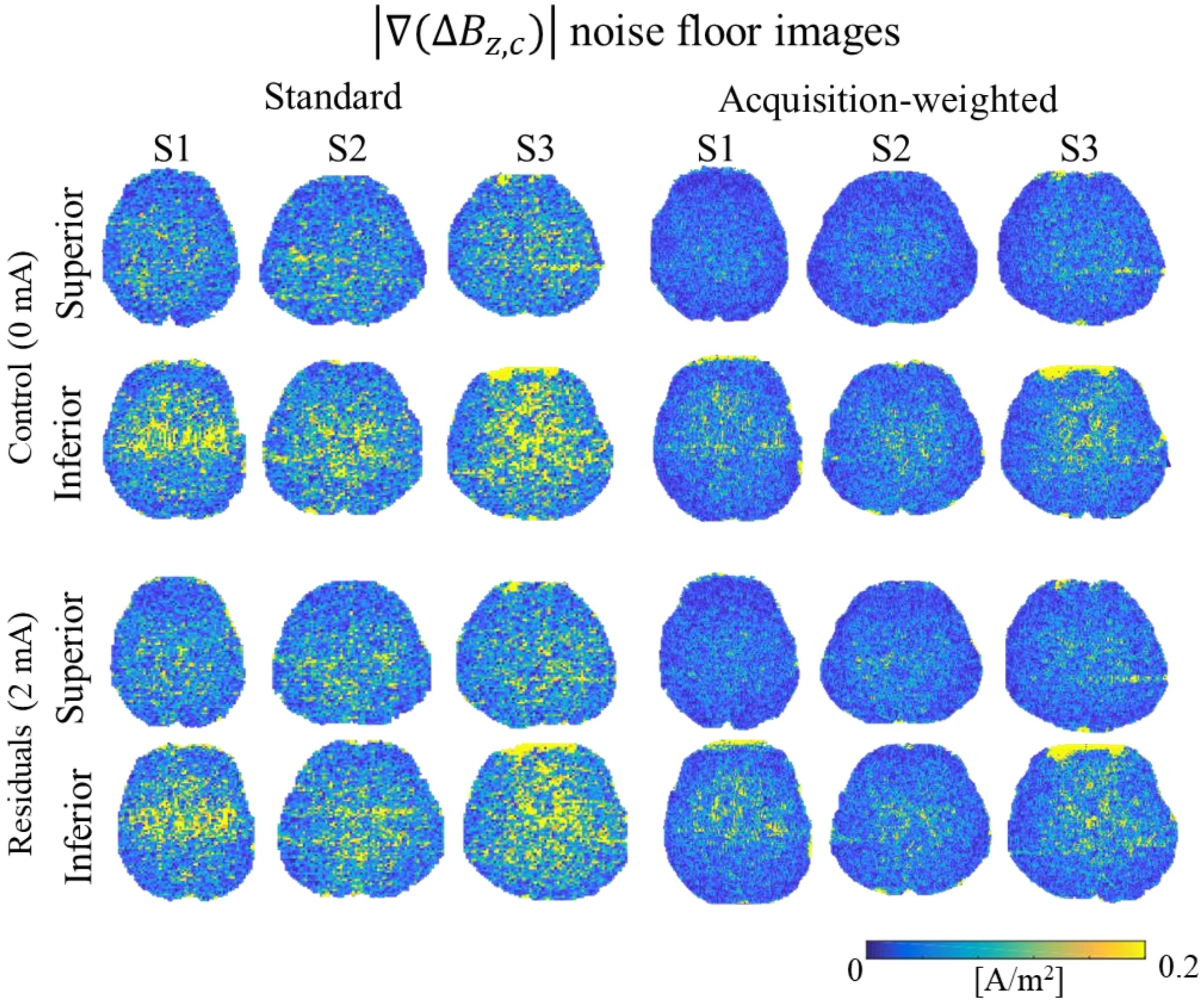


**Supplementary Figure S5.** Experiment 2: Standard vs. acquisition-weighted (AW) human in-vivo brain MRCDI. The results are shown for measurements of superior and inferior brain slices for all subjects. Images of the norm of the gradient of the current-induced magnetic field $|\nabla(\Delta B_{z,c})|$ are shown. Residual images (simulations subtracted from the magnetic fields measurements performed with LOOP-SETUP) are compared with control measurements without current flow, and similar noise floors are observed. Employing AW improves the resolution and the noise floors significantly compared to the standard acquisition scheme.

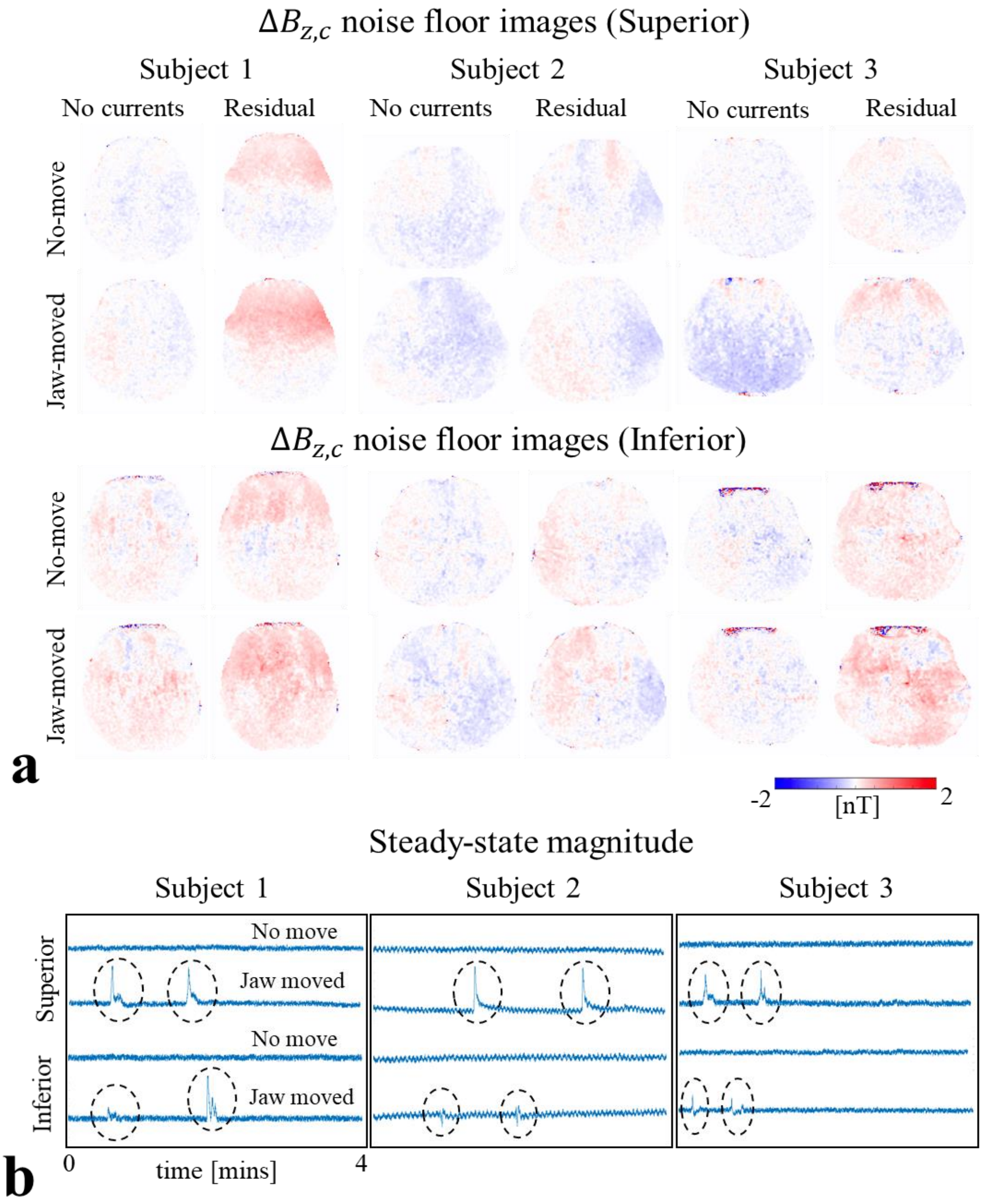


**Supplementary Figure S6.** Experiment 3: Navigators. The results are shown for measurements of upper and lower brain slices for all subjects. The measurements for two conditions are compared: The subject performs no intentional movement (No-move) versus intentional jaw movements twice during the scan (Jaw-moved). (a) Jaw movement-induced noise floor increases in the $\Delta B_{z,c}$ measurements are marginally observable. (b) The MR magnitude signal acquired from the navigator fluctuates during the two jaw movements, and stays constant for the rest of the scan (here we show only the control measurements without current flow, as the results were similar for the recordings with current flow). The results are consistent over subjects.

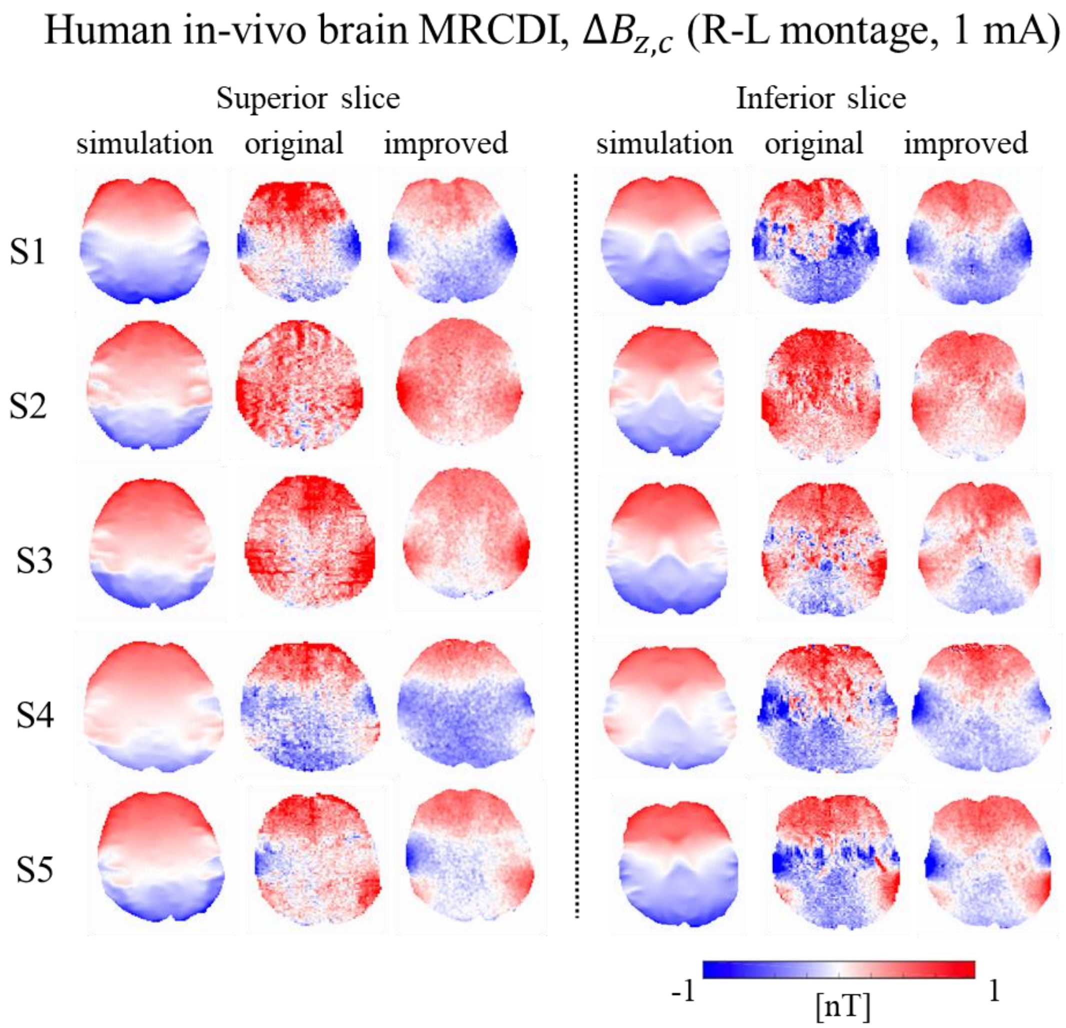


**Supplementary Figure S7.** Experiment 4: Human in-vivo brain MRCDI measurements and simulations for both the superior and inferior slices. The performance of our original method is directly compared with our improved method based on measurements with TES currents injected in anterior-posterior R-L direction. The measurements are consistent, but the improved method significantly reduces the $\Delta B_{z,c}$ noise floors, and enhances the image quality and resolution. Analogous to the A-P results, our improved method successfully resolves the artefacts near ventricles. The $\Delta B_{z,c}$ simulations and measurements for R-L are also in the same range, but show more differences in their spatial patterns when compared to the A-P results. The dependence of TES simulation accuracy on the injection configuration is beyond the scope of this study.

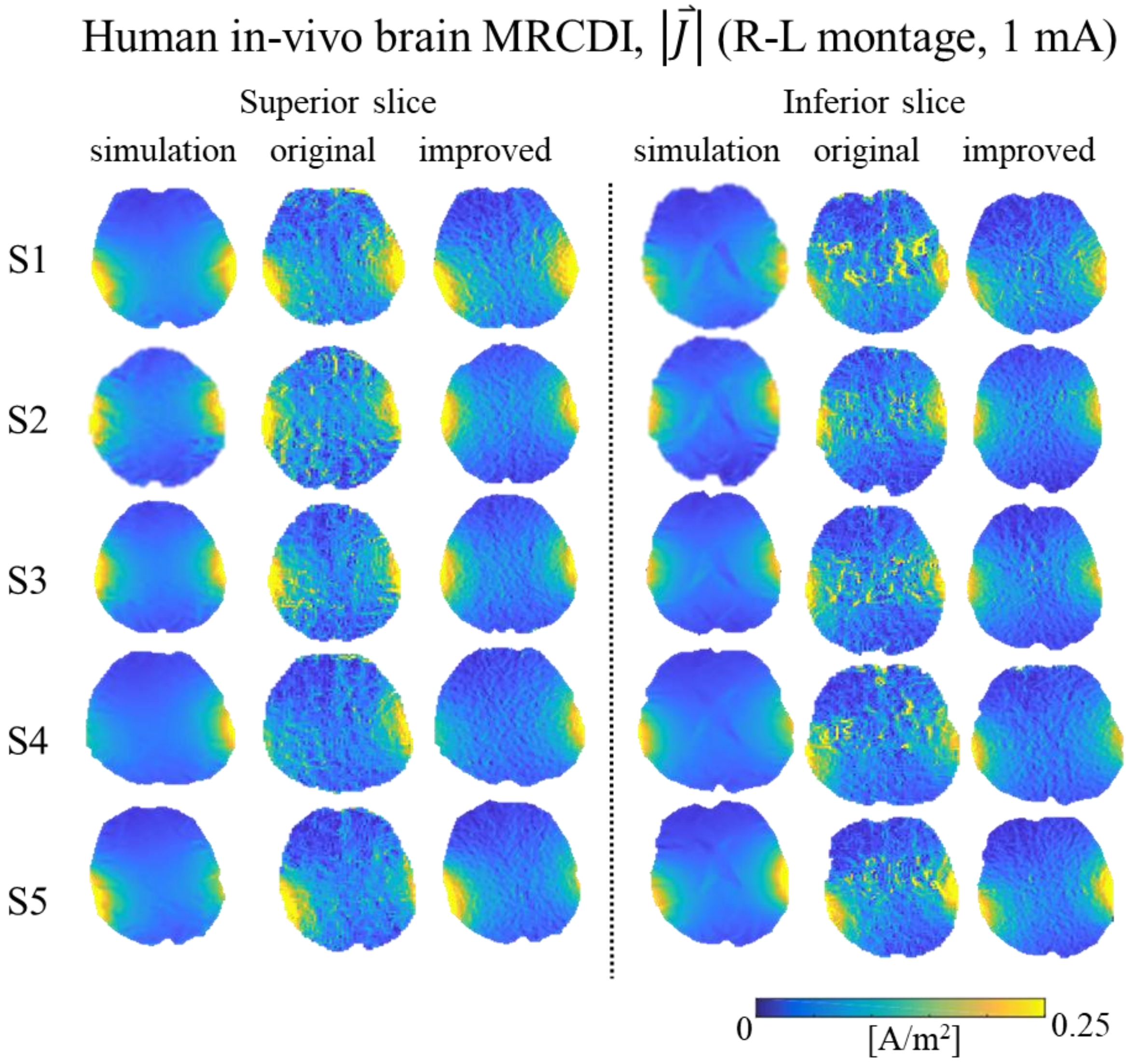


**Supplementary Figure S8.** Experiment 4: Human in-vivo brain MRCDI current flow reconstructions based on $\Delta B_{z,c}$ measurements and simulations for both the superior and inferior slices. The performance of our original method is directly compared with our improved method based on measurements with TES currents injected in anterior-posterior R-L direction. Our improved method greatly improves the noise floors and enhances the image quality and resolution. Analogous to the A-P results, our improved method successfully resolves the artefacts near ventricles. The reconstructed current flows from the simulations agree well with the measurements, but generally lack detailed spatial information. Compared to $\Delta B_{z,c}$, the simulated and measured current flow reconstructions show more similar distributions, which may also be caused by the approximations used in the projected current density reconstruction algorithm.
